# Quantile Regression for biomarkers in the UK Biobank

**DOI:** 10.1101/2023.06.05.543699

**Authors:** Chen Wang, Tianying Wang, Ying Wei, Hugues Aschard, Iuliana Ionita-Laza

## Abstract

Genome-wide association studies (GWAS) for biomarkers important for clinical phenotypes can lead to clinically relevant discoveries. GWAS for quantitative traits are based on simplified regression models modeling the conditional mean of a phenotype as a linear function of genotype. An alternative and easy to apply approach is quantile regression that naturally extends linear regression to the analysis of the entire conditional distribution of a phenotype of interest by modeling conditional quantiles within a regression framework. Quantile regression can be applied efficiently at biobank scale using standard statistical packages in much the same way as linear regression, while having some unique advantages such as identifying variants with heterogeneous effects across different quantiles, including non-additive effects and variants involved in gene-environment interactions; accommodating a wide range of phenotype distributions with invariance to trait transformation; and overall providing more detailed information about the underlying genotype-phenotype associations. Here, we demonstrate the value of quantile regression in the context of GWAS by applying it to 39 quantitative traits in the UK Biobank (*n* > 300, 000 individuals). Across these 39 traits we identify 7,297 significant loci, including 259 loci only detected by quantile regression. We show that quantile regression can help uncover replicable but unmodelled gene-environment interactions, and can provide additional key insights into poorly understood genotype-phenotype correlations for clinically relevant biomarkers at minimal additional cost.

## Introduction

GWAS are commonly used to identify genetic associations with phenotypes of interest, and have been successful in identifying a large number of associations with diverse traits and diseases. Although some of these genetic discoveries have informed successful therapeutic development^1^, the translational potential of GWAS discovery remains largely unrealized. Two reasons for the translational gulf separating GWAS discoveries from clinical application is the overall complex relationship between genotypes and clinical phenotypes, and the fact that the models used to connect genotypes with phenotypes do not capture this complexity. In particular, it is widely recognized that the influence of genetic variants on traits and diseases can be complex and dynamic^2, 3^. For example, gene expression changes with aging and varies across tissues and phenotypic contexts^2, 4–7^. Despite awareness of this complexity, the standard models used to interrogate the genome remain highly simplified, largely because the very large sample sizes required for genetic discovery preclude incorporating known complexities into the analytic framework (e.g. relevant environmental measures may not be available at this scale). The emergence of large-scale data resources such as biobanks linked to electronic medical records such as the UK Biobank (UKBB)^8^ make investigation of more complex genotype-phenotype associations possible. In particular, the use of biomarkers relevant to clinical phenotypes of interest is a promising avenue to clinically relevant genetic discoveries. The UKBB project collected and followed ≈500,000 individuals (aged between 40 and 69 at enrollment) across the UK since 2006. GWAS, whole-exome and whole-genome sequencing data are already (or will be soon) available on all individuals, together with a rich variety of health data, including physical measurements, biomarkers, physical activities, imaging, cognition assessments, and lifestyle.

GWAS are based on linear regression (LR) models including linear mixed effect models such as BOLT-LMM and fastGWA^9, 10^ that jointly model all variants to increase power over simple LR models. Linear regression models allow one to test whether genetic variants are associated with the mean of a phenotype distribution. However, the effect of a genetic variant can go beyond the mean, and can include the lower or upper parts of the phenotype distribution. For example, individuals with high body mass index (BMI) can be affected at a different extent by a certain genotype than average BMI individuals. However, LR with its focus on the central part of the phenotype distribution may not capture these more complex associations.

An alternative, natural extension of LR, admittedly rarely used in genetic studies of quantitative traits, is based on quantile regression (QR^11^). QR allows one to model any quantile of the phenotype distribution and can be used to measure how a genetic variant associates with various quantiles of a trait or equivalently how a genetic variant impacts different parts of the phenotype distribution, including mean, variance and higher moments. There are several examples in the genetics literature of such effects despite the very limited use of non-LR techniques in GWAS^12–17^. Almost all of these existing studies focus on detecting genetic variants that impact the phenotypic variance. Several known examples include an earlier study by Yang et al.^12^ that showed that a SNP in the *FTO* gene is associated with both the mean and the variance of BMI, as well as several variance quantitative trait loci (vQTL) studies^13–17^. Also, Wang et al.^16^ have shown that gene-environment effects can be inferred from genetic variants associated with the phenotypic variance without the need of measuring environmental factors. More generally, genetic associations can be complex due to underlying heterogeneity in the population and disease model, and the effects of gene-gene and gene-environment interactions^18, 19^. Similarly, non-additive effects including dominance and epistasis are not usually tested in GWAS because it is assumed that an additive model captures most of the contribution of a locus to a trait^20^. For eQTLs, several recent studies have shown the dynamic nature of eQTLs depending on the context or environment, providing evidence of widespread gene-environment interaction that can play a role in disease development^21^.

Along this direction, several methods exist to identify genetic variants that control phenotypic variance by testing for the homogeneity of variance including Levene’s test^22^, the Brown-Forsythe test^23^, the correlation least squared test^24^, a double generalized linear model test^25, 26^, and a Bayesian test^27^. However, mean and variance can only partially reveal heterogeneous effects. Recent applications of variance-based tests to biomarkers in UKBB revealed a small number of vQTLs, and 90% of those were already detected by LR^16, 17^. Different from testing moments, quantile regression based tests as emphasized here target distribution-wise differences, and can lead to detection of more general distributional differences. Even though differences at different quantiles can be related to moments, quantiles are easier to interpret and provide richer information about the detected associations.

The basic idea of QR is that genotypes might have different effects at different quantiles of the phenotype distribution. QR techniques are an active area of research in statistics and various efficient estimation algorithms and variable-selection methods have been developed for high-dimensional data^33–35^. These techniques are however not commonly utilized in genetics with a few exceptions^15, 36^, despite the similarity to LR and the availability of efficient implementations for genome-wide analysis in standard statistical packages^37, 38^. One likely reason is the difficulty in interpreting the results. For LR a regression coefficient tells how much the mean of the phenotype changes with 1-unit change in the predictor/genotype variable. For QR the interpretation is similar, namely a QR coefficient represents how much a specific quantile of the phenotype distribution changes with a 1-unit shift in the predictor/genotype distribution (e.g. for the 0.5 quantile, this would correspond to the change in median). This is indeed less intuitive, and an additional challenge for QR is that one gets quantile specific coefficients over a range of quantile levels. However, the upside is that we can obtain a more nuanced understanding and a better characterization of the underlying genotype-phenotype association, even at those associations identified by LR. In particular, it is possible that specific genetic variants only have an effect at upper or lower quantiles of a phenotype due, for example, to gene-environment interactions, and such quantile-specific effects would be masked in a vanilla LR method. Another great feature of QR is that it does not have assumptions on the error distribution, which means that it is robust to non-gaussian errors and outliers. While in linear regression traits are usually normalized using, for example, rank-based inverse normal transformation and results (type-1 error and power) can depend on the particular transformation^28^, with quantile regression results are invariant to monotonic transformations and therefore no particular trait transformation is required. That trait transformations should not affect our conclusions based on the regression results is of course highly desirable. Furthermore, that also helps when certain transformations are preferable to get a more natural interpretation of the association patterns.

The availability of rich phenotype and genotype data from the UKBB provides an opportunity to test the QR methodology in biobank scale data. Furthermore, the focus on biomarkers can lead to the identification of potentially more interesting associations for clinical phenotypes. In particular, it is often the case that high-risk subgroups (e.g. high risk for heart disease) have either high or low values for risk factors, so when modeling intermediate risk factors that are continuous, such as BMI or LDL cholesterol, using quantile regression can be very helpful as it can help identify variants with larger effects on subgroups of individuals but with modest or no contribution overall. Here, we investigate the performance of standard QR tools in the context of GWAS using 39 quantitative traits in the UKBB. In particular, we consider a series of quantile levels and use a rank-score approach^29^ to identify variants with impact on the distribution of each trait. We further test the detected QR associations for gene-environment interaction effects.

## Results

### Overview of linear and quantile regression techniques

We describe here briefly the main statistical models we compare in our analyses. We consider first the linear additive model normally used in GWAS and then discuss the quantile regression model we propose as a complementary strategy. More details are in the Methods section.

#### Linear Regression (LR)

We assume that we have *n* independent samples from a population, and for the *i*th subject, we denote by *Y_i_* a continuous (quantitative) phenotype, by *X_i_* = (*X_i_*_1_, *X_ip_*)′ the genotypes at *p* genetic variants, and by *C_i_* = (*C_i_*_1_, *C_iq_*)′ covariates we want to adjust for, such as age, gender and principal components of genetic variation. Furthermore, we denote by *Y* = (*Y*_1_, … , *Y_n_*)′ the *n* 1 sample phenotype vector, by *X* the *n × p* genotype matrix, and by *C* the *n × q* matrix for covariates. We assume that *Y* is normalized. In GWAS, the typical approach is to perform marginal (unconditional) testing for each variant *j* at a time. Therefore the model is:

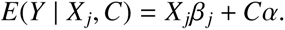

We define the error projection matrix *P_C_* = *I* - *C*(*C*′*C*)^−1^*C*′. We project *Y* and *X* onto the orthogonal complement of the column space of *C*, then we get

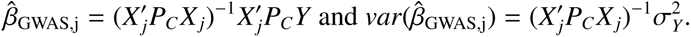

We denote by *β̂*_GWAS_ = (*β̂*_GWAS,1_, · · · , *β̂*_GWAS,p_)′ and the vector of marginal *z* scores as

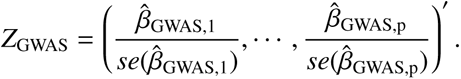

#### Quantile Regression (QR)

We denote the *τ*th conditional quantile function of *Y* as *Q_Y_* (*τ X*, *C*). Then in linear quantile regression we can write the conditional quantile regression model for *j*th variant and given a specific quantile level *τ* ∈ (0, 1) as:

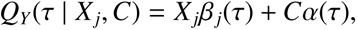

where *β _j_*(*τ*), and α(*τ*) are quantile-specific coefficients and can be estimated by minimizing the pinball loss function^11^. For median regression (*τ* = 0.5), that amounts to finding the regression line that minimizes the sum of the absolute values of the residuals (same as in LAD - least absolute deviations) rather than the sum of the squared residuals as in linear regression. More generally, the quantile-specific coefficients are estimated by minimizing a weighted sum of absolute residuals which can be formulated as a linear programming problem that can be solved efficiently. For quantile regression, a commonly used hypothesis testing tool is the rank score test^30, 31^. The rank score test statistic for a fixed quantile level *τ* is defined as:

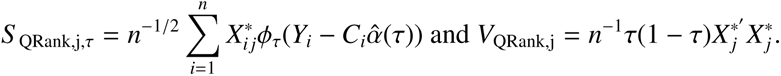

where *X*^∗^ = *P_C_ X*, and φ_*τ*_(*u*) = *τ* − *I*(*u* < 0) is the *τ*-pinball loss. Under the null hypothesis 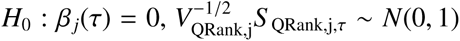. We note that the asymptotic distribution of the test statistics is independent of the particular distribution of the phenotype. Hence it can be applied to any phenotype without requiring a pre-transformation to achieve normality.

We denote by 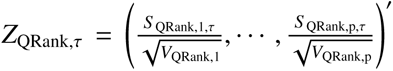 for a specific quantile level *τ* ∈ (0, 1).

Throughout this paper, in both simulations and real data analyses, we use 9 quantiles spaced equally at 0.1 intervals. We combine quantile specific p-values from the marginal tests via Cauchy combination^32^.

### Evaluation of QR in simulations

We use UKBB genotype data on 20,000 white British unrelated individuals, and focus on one 1MB region (chr8:19,291,998-20,291,997; GRCh37/hg19) with 4,148 variants. We assume only one causal variant selected at random, and simulate phenotype data using several models, including a homogeneous linear model (with effects being constant across quantiles) with the error distribution being *N*(0, 1) or a heavy tail *Cauchy*(1, 0) distribution; a local model where effects only exist at the upper quantiles; and a model with both additive and dominance effects. Trait is normalized using rank-based inverse normal transformation for LR only; raw trait values are used for QR.

#### Homogeneous linear model

We assume that the phenotype *Y* follows the model:

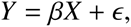

where *X* is the genotype at the causal variant, *ϵ N*(0, 1). In this model, the quantile effect is constant across all quantiles (Fig. 1(a)). We then assess the power of LR and QR based on 100 Monte Carlo replicates when varying the effect size *β*: 0.01 < *β* < 0.2; the causal variant is chosen randomly for each replication, with MAF > 1%. We show that overall LR has a slight power increase over QR under this scenario with homogeneous association and normal error (Fig. 1(a)).

**Figure 1:**
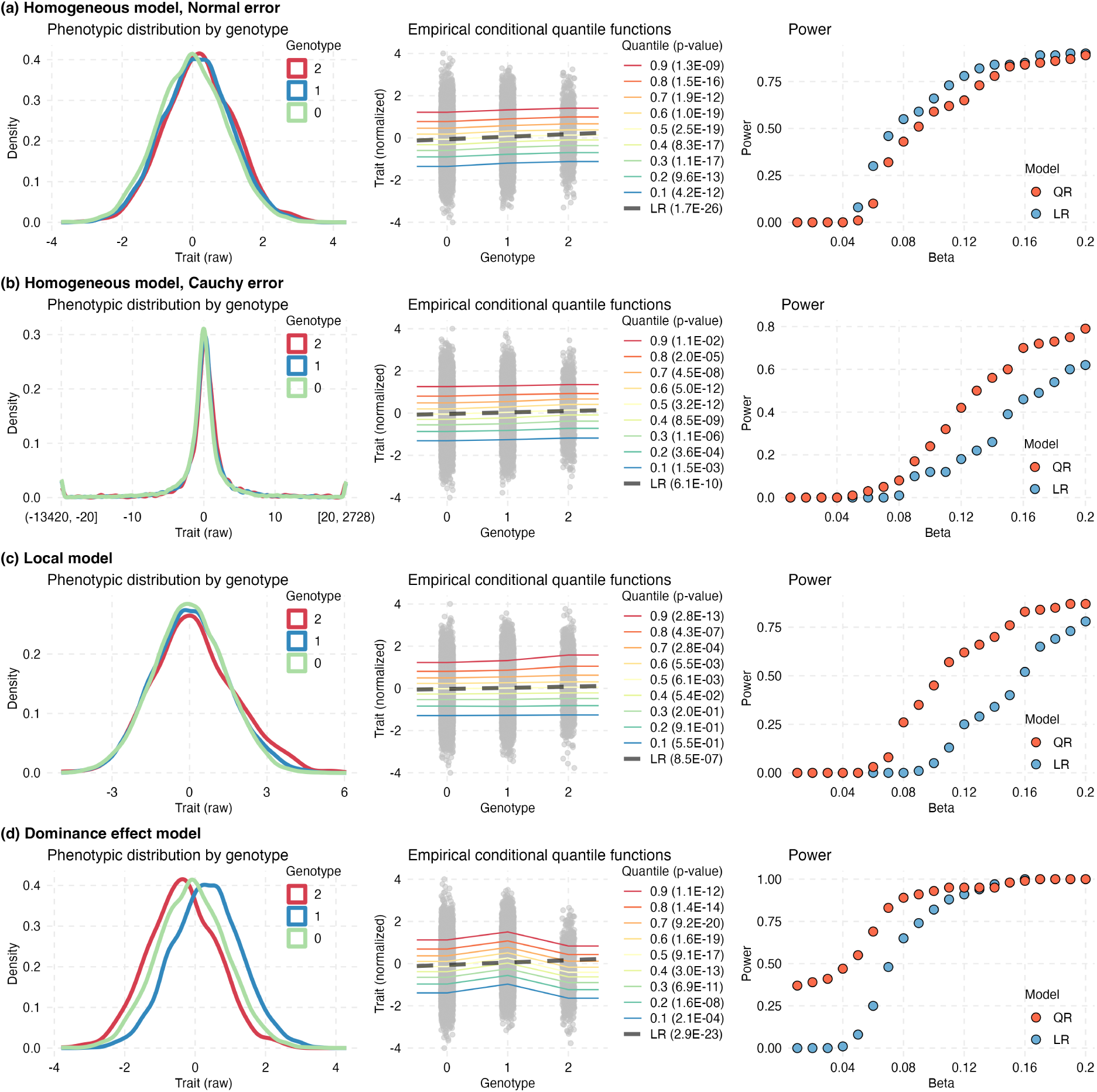
Power results in simulations for (a) homogeneous model, Normal error distribution, (b) homogeneous model, Cauchy error distribution, (c) local model, and (d) model with additive and dominance effects. In the homogeneous model, effects are additive and constant across quantiles, error has a Gaussian or Cauchy distribution. In the local model, effects are only present for quantile level *τ* > 0.7. Power is shown when *beta* varies from 0.01 to 0.2. For the model with additive and dominance effects, the dominance effect is fixed at 0.2 while the additive effect varies from 0.01 to 0.2. The figures on the left show the densities of the raw trait values by genotype group for one replicate. The figures in the middle show the empirical conditional quantile functions for *τ* = 0.1, 0.2, 0.3, 0.4, 0.5, 0.6, 0.7, 0.8, 0.9 for the same replicate, under the homogeneous and local models for *beta* = 0.1, and *beta_A_*= 0.1 and *beta_D_* = 0.2 for the additive+dominance model. Shown in black is also the LR line fitted to the data.

#### Model with Cauchy errors

We assume that the phenotype *Y* follows the model:

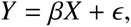

where *X* is the genotype at the causal variant, *ϵ Cauchy*(1, 0). In this model, quantile effects are constant across quantile levels, but the trait has a heavy tail. QR has higher power than LR in this setting (Fig. 1(b)).

### Local model with effects only at upper quantiles

We additionally simulate data under a local model with effects only at the upper quantiles: *τ* ∈ [0.7, 1] (Fig. 1(b)). Such a model can correspond to a gene-environment interaction, where, for example, a genetic variant only shows effect on a phenotype in a certain environment.

Specifically, we assume that the conditional quantile function of the phenotype *Y* can be written as:

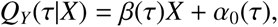

where *β*(*τ*) = 5*β*(*τ* − 0.7)/(1 − 0.7) when *τ* > 0.7 and *β*(*τ*) = 0 otherwise. We can view α_0_(*τ*) as the quantile function of *Y* when *X* = 0. The error distribution (α_0_^−1^(*τ*)) is assumed to be normal. To simulate *Y* from this model, we use the inverse quantile approach, where we randomly draw a *U*(0, 1) random variable as *τ*, and plug it into the conditional quantile function *Q_Y_* (*τ X*).

Under this local model, QR can have significant increase in power over LR (Fig. 1(c)).

#### Model with both additive and dominance effects

Finally, we consider a model that includes both additive and dominance effects as follows:

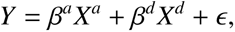

where *X^a^* corresponds to the additive coding 0, 1 and 2, while *X^d^* is coded as *p*/(1 *p*), 1, (1 *p*)/*p*) with *p* being the minor allele frequency (MAF), and *ϵ N*(0, 1). When the dominance effect is strong enough, QR can indeed be more powerful (Fig. 1(d), S1).

### Applications to UKBB GWAS for 39 quantitative traits

#### GWAS discoveries by LR and QR for 39 traits

To evaluate QR and contrast the results with LR, we use the UKBB data (between 264,854 and 325,825 white British, unrelated individuals and ~ 8.6 million SNPs), and perform GWAS for 39 quantitative traits (Fig. 2). We adjust for several covariates including age, sex, batch, and 10 principal components (PC) of genetic variation. We identify significant SNPs using a genome-wide significance threshold of 5×10^−8^ for each trait (Fig. S2-S11). We define a locus as a 1Mb segment around the top p-value SNP in a region. LR detects a larger number of significant associations relative to QR (2,039,518 genome-wide significant SNPs and 9,964 loci vs. 1,517,669 SNPs and 7,297 loci). Most of the QR discoveries are shared with LR, but QR also identifies new loci that LR misses (Fig. 2 and 3(a)). Loci detected by QR but missed by LR (i.e. QR-LR), and loci detected by LR but missed by QR (i.e. LR-QR) tend to have less significant p-values relative to loci detected by both (i.e. LR&QR), suggesting more heterogeneous or lower effect sizes (Fig. S12).

**Figure 2:**
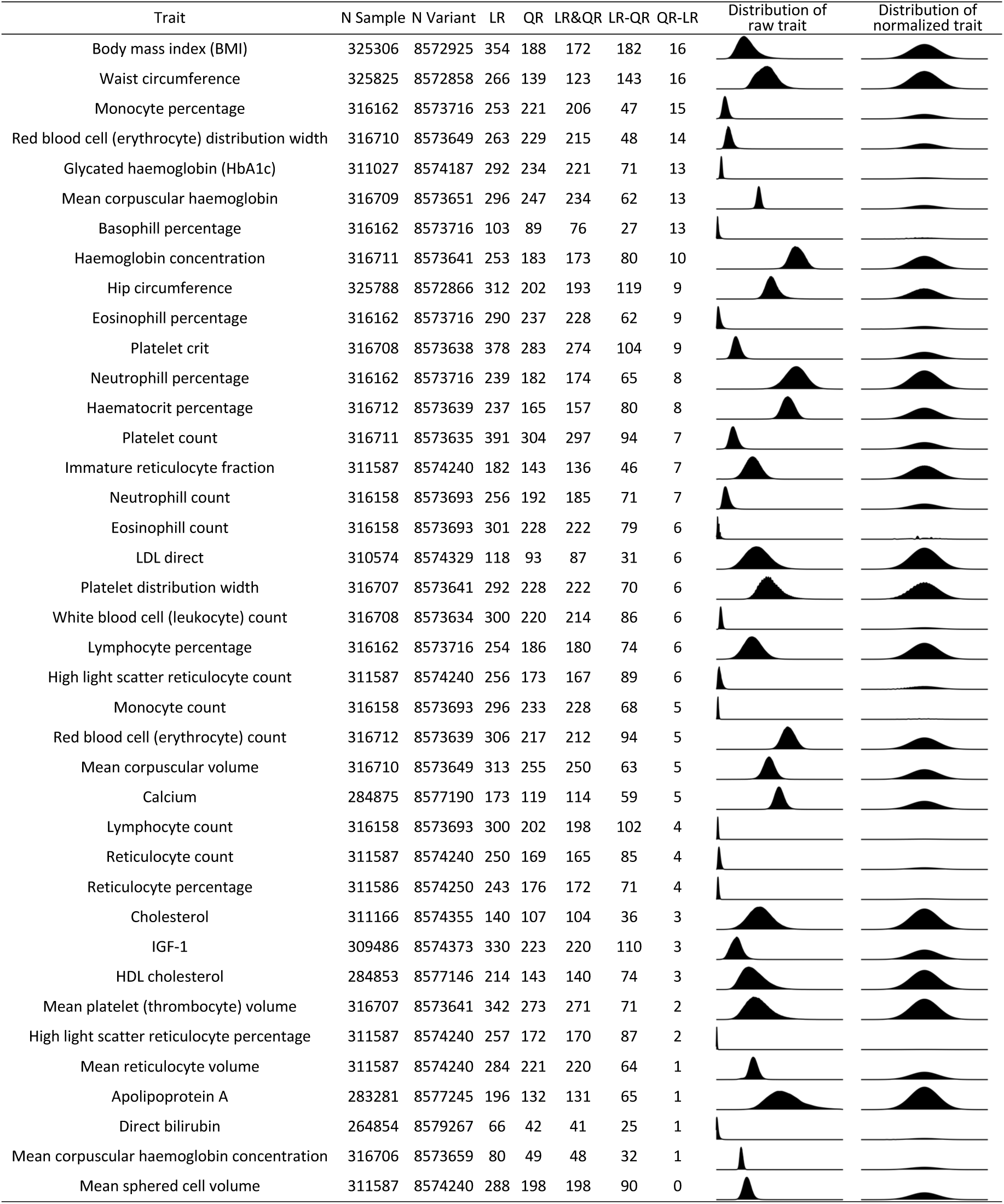
Number of genome-wide significant loci identified by LR and QR for each of 39 traits. LR-QR are 1 Mb loci identified by LR but missed by QR; QR-LR are 1 Mb loci identified by QR but missed by LR; LR& QR are 1 Mb loci identified by both LR and QR. For each trait we also show the distribution of the raw trait, and after rank-based inverse normal transformation.

**Figure 3:**
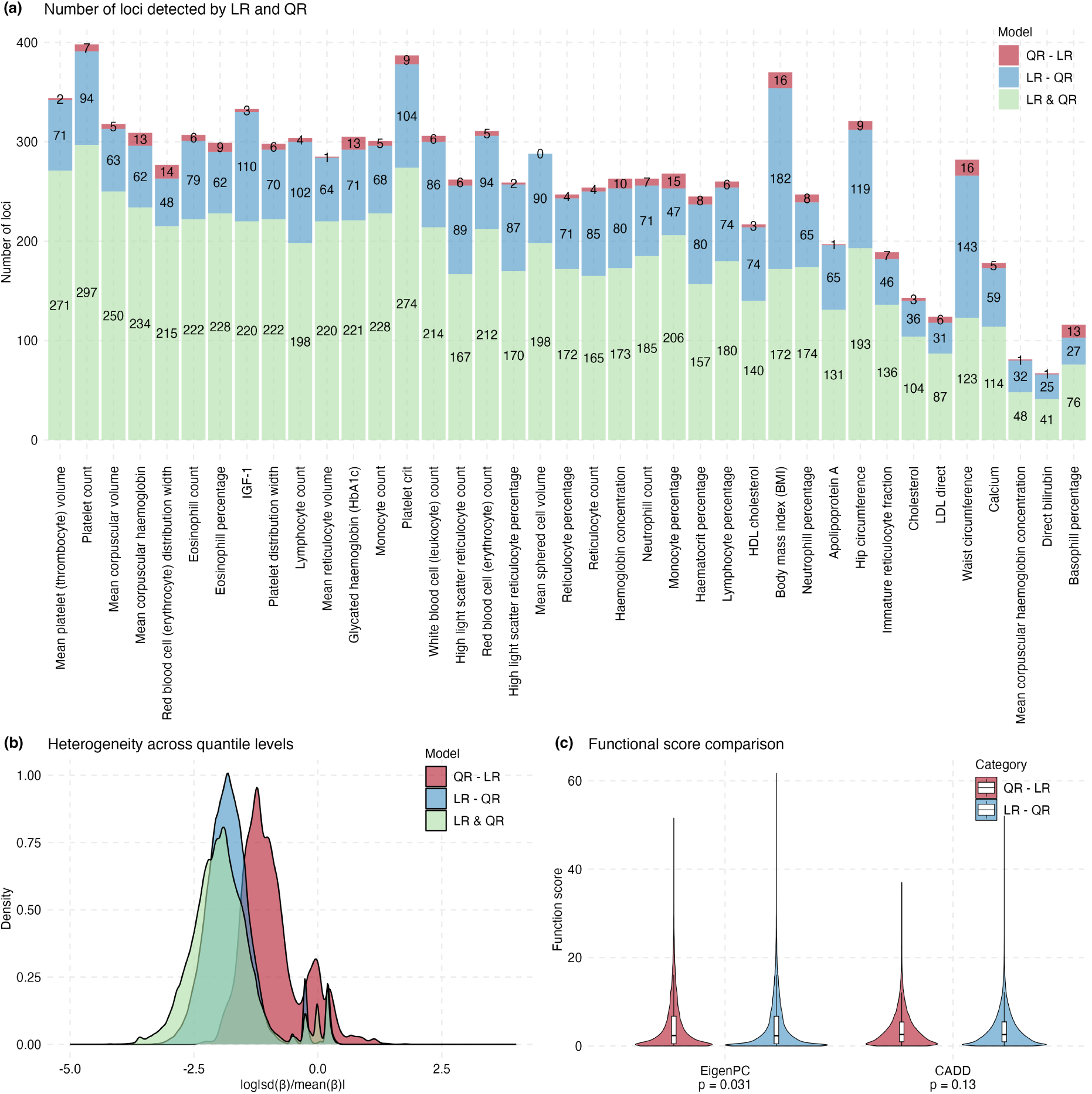
Number of genome-wide significant loci identified by LR and QR for each of 39 traits, effect size heterogeneity, and functional score comparison. (a) Number of genomewide significant loci. Stacked barplots show the corresponding number of significant loci for LR&QR, LR-QR and QR-LR. (b) Densities of heterogeneity indexes for GWAS associations for 39 traits. Densities are shown for three sets of associated SNPs: LR&QR, LR-QR and QR- LR. (c) Functional scores (EigenPC and CADD) for the associated SNPs detected by QR-LR compared to those from LR - QR. One-sided p-values (QR-LR>LR-QR) from the Kolmogorov- Smirnov test are also shown comparing the functional effects for variants identified by QR-LR vs. LR-QR.

#### Replications of QR and LR associations

We attempted to replicate the genome-wide significant associations above in an independent dataset of European individuals from UKBB (*n* ≈ 34, 000 39, 000). Given the much smaller sample sizes for the replication datasets relative to the discovery datasets, we report replication at nominal significance level (p-value < 0.05). One challenge with QR is that the result is a composite of results across multiple quantile levels and the p-value is obtained by Cauchy combination of individual quantile-level p-values which makes it difficult to take the sign of the associations into account when considering replication. With this caveat in mind, we noticed that QR has replication rates at least as high as LR (67% vs. 61% on average across the 39 traits, Fig. S13, Table S1). For LR, sign consistency tends to be very high (≈ 96%).

#### QR-LR vs. LR-QR

Since QR allows us to estimate quantile specific effects across a whole range of quantiles, we can quantify the degree of heterogeneity in effects across the quantiles for individual SNPs. Specifically, for a SNP we compute a heterogeneity index as the log transformed ratio between the standard deviation and the absolute mean of the estimated quantile coefficients *β*(*τ*)’s: log sd(*β*(*τ*))/mean(*β*(*τ*)) . The higher the number, the more heterogeneity in the quantile effects. As expected, SNPs identified by QR but missed by LR (QR p-value < 5*e* 08, LR p-value > 5*e* 08) are the most heterogeneous (Fig. 3(b)). Furthermore, we use several functional annotation scores in the literature, including Eigen-PC^39^ and CADD^40^, to compare functional scores for variants identified by QR but missed by LR (QR p-value < 5*e* 08, LR p-value > 5*e* 08) vs. variants identified by LR but missed by QR (LR p-value < 5*e* 08, QR p-value > 5*e* 08). Variants identified by QR but missed by LR have significantly higher functional scores relative to those only identified by LR (Eigen-PC p=0.031, Fig. 3(c)).

#### BMI and LDL

Next we focus on two traits, BMI and LDL, to provide more specific results. For BMI, QR identifies 188 loci, of which 172 are shared with LR and 16 are unique to QR (Fig. 4(a)-S14, Table 1). Variants identified by QR but missed by LR have heterogeneous effects across quantile levels as expected, with larger effects at the upper quantiles (Fig. S15). For LDL, QR identifies 93 loci, with 87 of them being shared with LR, and 6 being unique to QR (Fig. 4(b)-S16, Table 1). We noticed a similar pattern as with BMI, with variants identified by QR but missed by LR having larger effects at the upper quantiles (Fig. S15). Note that, in general, we expect associations to be equally likely to be linked to the upper and lower quantiles (Supplemental Figure 1). Moreover, variants identified by QR but missed by LR have more heterogeneity (Fig. S17(a)) and higher functional scores (Fig. S17(b)) relative to those only identified by LR.

**Figure 4:**
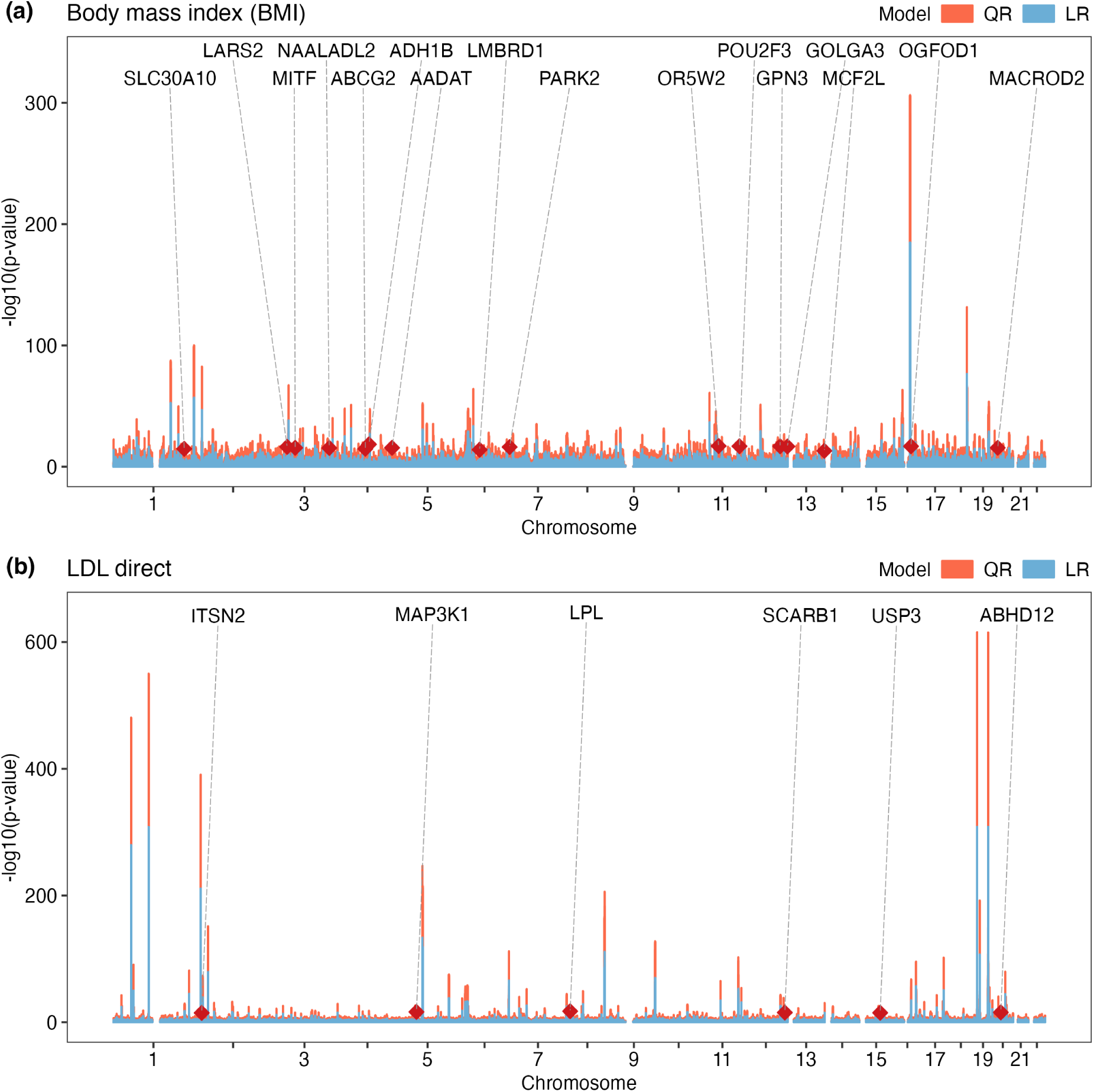
Manhattan Sunset plot for BMI and LDL. -log10(p-value) of the significant associations (p-value < 5 10^−8^) detected by the LR (blue bars) and QR (red bars) for (a) BMI and (b) LDL are plotted against the genomic positions. The lead QR association at a locus detected by QR - LR is represented by a red diamond with the name of the closest protein-coding gene indicated.

**Table 1:** Lead QR SNPs at QR-LR significant loci in BMI and LDL. Positions (POS) of the variants are in GRCh37 coordinates. Effect allele frequency (EAF) is counted based on the effect allele (A1) from GWAS. INFO refers to the imputation quality score. LR, QR and quantile specific p-values are also shown.

| Trait | Locus | SNP | CHR | POS | A1 | A2 | EAF | INFO | LR | QR | Quantile level p-value |  |  |  |  |
| --- | --- | --- | --- | --- | --- | --- | --- | --- | --- | --- | --- | --- | --- | --- | --- |
|  |  |  |  |  |  |  |  |  |  |  | QR 0.1 | QR 0.3 | QR 0.5 | QR 0.7 | QR 0.9 |
| BMI | SLC30A10 | rs12037905 | 1 | 219628036 | T | C | 0.418 | 0.995 | 2.35E-05 | 8.16E-09 | 5.63E-01 | 6.59E-02 | 6.08E-04 | 2.75E-07 | 4.41E-07 |
|  | LARS2 | rs9852062 | 3 | 45373442 | T | A | 0.444 | 0.988 | 3.63E-07 | 1.25E-08 | 3.67E-01 | 1.45E-05 | 5.63E-08 | 9.38E-08 | 8.82E-04 |
|  | MITF | rs9814379 | 3 | 70180381 | G | A | 0.077 | 0.988 | 3.42E-06 | 8.05E-09 | 8.97E-10 | 2.34E-05 | 2.70E-03 | 3.40E-02 | 2.43E-01 |
|  | NAALADL2 | rs4435653 | 3 | 175600730 | G | T | 0.311 | 0.995 | 6.80E-06 | 5.38E-09 | 5.67E-02 | 1.04E-02 | 1.40E-03 | 5.91E-04 | 5.98E-10 |
|  | ABCG2 | rs4148155 | 4 | 89054667 | G | A | 0.114 | 1.000 | 8.26E-06 | 2.08E-08 | 3.35E-01 | 1.14E-03 | 2.31E-09 | 3.19E-05 | 1.13E-02 |
|  | ADH1B | rs1229984 | 4 | 100239319 | T | C | 0.022 | 1.000 | 5.36E-08 | 7.07E-10 | 9.04E-02 | 1.68E-03 | 3.82E-08 | 1.05E-10 | 3.01E-05 |
|  | AADAT | rs55920177 | 4 | 171609715 | T | A | 0.128 | 0.994 | 2.11E-06 | 1.26E-08 | 1.79E-01 | 1.31E-03 | 3.89E-07 | 1.42E-05 | 6.92E-03 |
|  | LMBRD1 | rs72906423 | 6 | 70261968 | T | C | 0.078 | 0.952 | 2.74E-05 | 3.82E-08 | 5.71E-01 | 8.21E-02 | 9.71E-04 | 1.14E-06 | 4.31E-09 |
|  | PARK2 | rs68045643 | 6 | 163029709 | G | A | 0.138 | 0.996 | 2.11E-07 | 2.96E-08 | 1.89E-04 | 1.96E-05 | 5.19E-05 | 3.62E-04 | 3.20E-03 |
|  | OR5W2 | rs12363672 | 11 | 55684028 | C | A | 0.026 | 1.000 | 1.77E-07 | 4.99E-09 | 1.73E-03 | 2.39E-09 | 6.25E-06 | 3.34E-04 | 6.21E-02 |
|  | POU2F3 | rs7121784 | 11 | 120159052 | A | C | 0.160 | 0.998 | 5.15E-08 | 2.71E-08 | 6.94E-04 | 6.74E-02 | 3.87E-03 | 7.58E-07 | 6.12E-06 |
|  | GPN3 | rs6606686 | 12 | 110903380 | G | C | 0.321 | 0.999 | 6.43E-08 | 2.35E-08 | 1.43E-02 | 2.67E-05 | 7.68E-09 | 2.75E-05 | 2.71E-03 |
|  | GOLGA3 | rs200089405 | 12 | 133402839 | AG | A | 0.311 | 0.989 | 3.89E-07 | 5.48E-09 | 1.03E-01 | 2.53E-03 | 8.04E-04 | 3.67E-06 | 6.09E-10 |
|  | MCF2L | rs35515197 | 13 | 113638124 | A | G | 0.028 | 0.963 | 3.10E-04 | 2.75E-08 | 5.76E-01 | 1.25E-02 | 1.10E-06 | 1.77E-06 | 6.71E-01 |
|  | OGFOD1 | rs78231432 | 16 | 56488440 | C | T | 0.132 | 0.997 | 2.33E-07 | 6.41E-09 | 1.47E-01 | 1.69E-03 | 2.05E-04 | 7.13E-10 | 1.72E-04 |
|  | MACROD2 | rs16996657 | 20 | 15816236 | C | T | 0.128 | 0.991 | 1.46E-06 | 1.85E-08 | 7.22E-02 | 1.42E-03 | 2.48E-06 | 8.82E-08 | 1.40E-02 |
| LDL | ITSN2 | rs78711174 | 2 | 24496531 | A | G | 0.269 | 1.000 | 7.50E-06 | 2.27E-08 | 2.35E-02 | 1.31E-01 | 1.28E-04 | 1.44E-05 | 7.56E-03 |
|  | MAP3K1 | rs9687846 | 5 | 55861894 | A | G | 0.200 | 1.000 | 1.35E-06 | 3.83E-09 | 5.80E-01 | 9.27E-03 | 6.89E-05 | 4.70E-10 | 4.84E-08 |
|  | LPL | rs326 | 8 | 19819439 | G | A | 0.292 | 1.000 | 1.06E-05 | 3.89E-11 | 1.80E-01 | 6.26E-02 | 7.61E-06 | 4.35E-12 | 1.42E-08 |
|  | SCARB1 | rs10846744 | 12 | 125312425 | C | G | 0.141 | 1.000 | 3.94E-06 | 1.18E-08 | 1.45E-01 | 8.16E-02 | 6.27E-06 | 4.02E-08 | 1.23E-07 |
|  | USP3 | rs56369308 | 15 | 63792257 | C | T | 0.342 | 0.998 | 3.24E-06 | 3.47E-08 | 4.55E-01 | 3.04E-02 | 3.94E-05 | 1.64E-08 | 5.93E-06 |
|  | ABHD12 | rs776504228 | 20 | 25362262 | C | CT | 0.443 | 0.987 | 7.46E-06 | 3.65E-09 | 6.38E-02 | 4.41E-03 | 4.52E-05 | 6.20E-08 | 7.08E-04 |

#### Some examples of QR-LR significant loci in BMI and LDL

We show here two specific examples for QR-LR loci for BMI and LDL. We focus on the two loci with the most heterogeneous SNPs. Both lead QR SNPs have protective effects against high BMI and high LDL, respectively. Namely, for BMI we show the locus corresponding to the lead QR SNP rs12037905 (Fig. 5(a)) which falls within LYPLAL1-antisense RNA1 (LYPLAL1-AS1), a long noncoding RNA gene which was found to be associated with BMI^41^ and lipid traits such as high-density lipoprotein (HDL) cholesterol and triglycerides (TG) levels^42^. Yang et al. described a mechanisms by which long noncoding RNA regulates adipogenic differentiation of human mesenchymal stem cells and suggested that LYPLAL1-AS1 may serve as a novel therapeutic target for preventing and combating diseases related to abnormal adipogenesis, such as obesity^43^.

**Figure 5:**
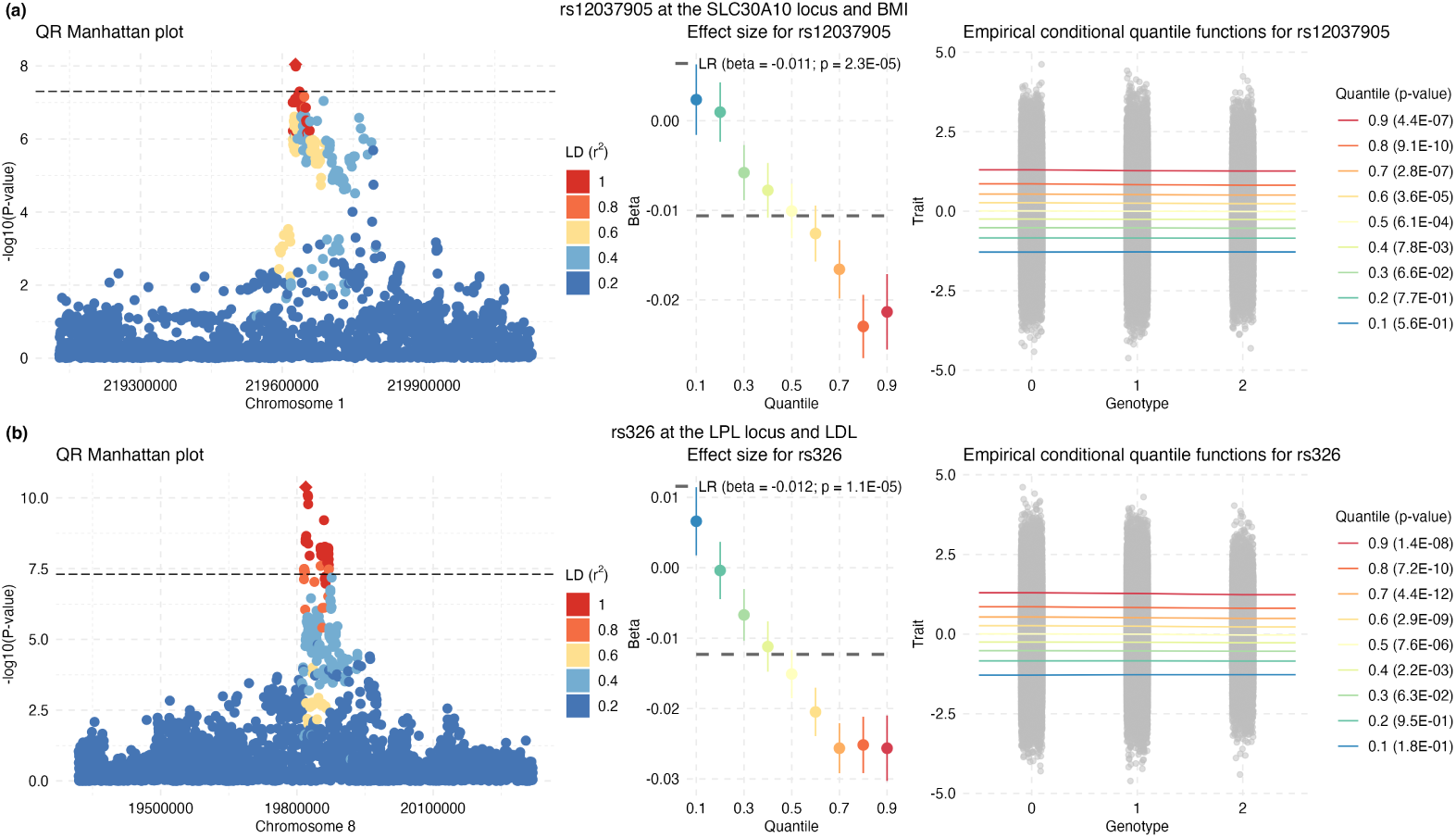
Examples of QR-LR significant loci in BMI and LDL. The Manhattan plots show -log10(p-value) of QR with the lead QR SNP per locus shown as diamond. Colors indicate the LD (*r*^2^) to the lead SNP for each variant. For the lead QR SNP at each locus, the estimated quantile specific effects SE are shown (middle panel), and the empirical conditional quantile functions for *τ* = 0.1, 0.2, 0.3, 0.4, 0.5, 0.6, 0.7, 0.8, 0.9 quantiles of BMI and LDL by genotype group (one gray dot per individual) are shown in the right panels.

For LDL, we show the lipoprotein lipase (*LPL*) with lead QR SNP rs326 (Fig. 5(b)) which is an intron variant within *LPL*. LPL, encoding lipoprotein lipase, plays an important role in the clearance of TG and maturation of HDL. Higher concentration of serum LPL is associated with a lower/higher level of TG/HDL cholesterol and contributes to a reduced risk of coronary artery disease (CAD)^44, 45^. rs326 at LPL is a pleiotropic variant associated with CAD^46^ and lipid metabolism^47^. Large lipid genetic studies have shown that the rs326 G allele is associated with increased HDL cholesterol levels and decreased TG, suggesting a protective effect for CAD risk^47^. In our QR analyses, the rs326 G allele is significantly associated with decreased LDL at upper quantiles (Fig. 5(b)), suggesting a protective effect against CAD risk among the individuals with higher levels of LDL.

#### Evidence of gene-environment interactions

Heterogeneous associations identified by QR may be the result of gene-environment interactions. Even though gene-environment interactions are believed to be common, detecting them is challenging due to several factors including lower statistical power relative to detecting main effects, low prior probability of specific gene-environment interactions, and sensitivity of results to the scale etc. Furthermore, the relevant environmental exposures can be missing from the study, and even when measured, accurate measurements of exposure may not be available. Here we employ a two-stage design, where we first identify significant SNPs from QR or LR, and then test for interactions between these SNPs and particular environmental factors using Bonferroni method to adjust for the number of gene-environment tests in the second stage (note that the two stages are independent^48^).

We selected five environmental factors (same as in^16^) that are available in the UKBB, including sex, age, physical activity (PA), sedentary behavior (SB) and smoking, and tested for interactions with top SNPs detected by QR using the following model:

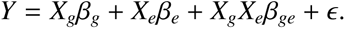

We tested *β_ge_* = 0 using a standard ANOVA test and adjusted for multiple testing using Bonferroni method.

Overall, using lead QR SNPs at 7,297 loci across 39 traits we identify 139 significant gene-environment interactions (significant after multiple testing adjustment for tests for each phenotype), 58 of which are significant at the study-wide level (*p* < 0.05/(7, 297 5) = 1.37*e* 06), after adjusting for all gene-environment tests across all phenotypes (Table S2 and Fig. 6(a)). To test the enrichment of gene-environment interaction effects among the lead QR variants, we repeated the gene-environment tests using the same number of randomly selected significant variants (LR or QR) across 39 traits. Over 1,000 replicates, we observed, on average, 16 significant gene-environment interaction effects, significantly lower than the 58 observed interactions with leading QR variants (Fig. 6(b)). If we focus on QR-LR loci, only 2 study-wide significant interactions are detected among 259 × 5 tests: *ARHGEF3* with age for platelet distribution width (*p* = 9.09*e* 16), and *RSPO3* with sex for waist circumference (*p* = 7.49*e* 07). Note that this is still a 11-fold enrichment over the 2 study-wide significant interactions detected at the LR-QR loci among 2, 926 5 tests (*p* = 0.03). Even though the loci detected by QR-LR and LR-QR are both weaker than loci detected by LR&QR (Fig. S12), variants at loci detected by QR-LR seem to be more functionally relevant than those at loci detected by LR-QR. Similar results hold if we restrict to BMI and LDL (Fig. S18), for which most of the significant interactions are with age or sex. For example, for LDL we detected significant interactions between variants in *PCSK9, APOB, CELSR2, LPA* and age. For BMI, we identified significant interactions between a variant in *FTO* and physical activity, and a variant in *FAIM2* and sedentary behavior.

**Figure 6:**
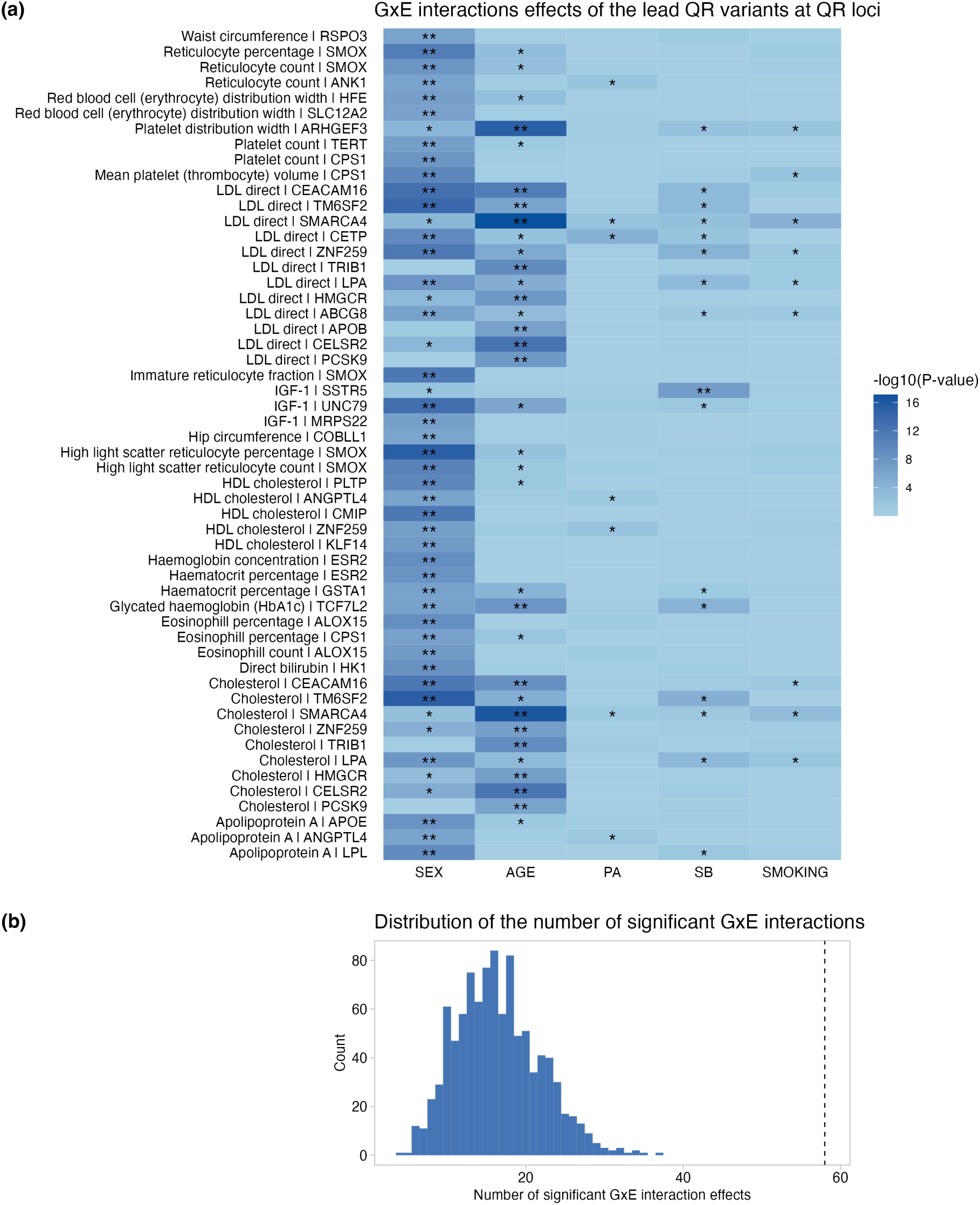
Significant gene-by-environment interaction effects of the lead QR variants for 39 traits. (a) The heatmap shows -log10(p-value) of gene-environment interaction tests for the lead QR association at the loci detected by QR. ”*” denotes nominally significant GxE interaction effects (p-value < 0.05) and ”**” denotes significant effects after Bonferroni correction (*p* < 0.05/(5 7, 297) = 1.37*e* 06). (e) The empirical distribution of the number of significant gene-environment interactions when variants are chosen randomly among significant variants identified by LR or QR.

#### Interaction of *ARHGEF3* with age for platelet distribution width

As mentioned above, one highly significant interaction was detected with rs1354034, a SNP with heterogeneous quantile effects that was completely missed by LR (Fig. 7). Interestingly, a strong non-additive association was previously observed for platelet width distribution at the same variant rs1354034, associated with increased expression of *ARHGEF3* in platelets, the association being highly specific to platelets and not detected in other major blood cell types^20, 49^. Platelet width distribution describes how similar the platelets are in size, with high values meaning great variation is size and being associated with vascular (blood vessel) disease or certain cancers. The interaction of rs1354034 with age may be indicative of increased platelet activation with age which leads to more variable platelet width^50^. Specifically, at younger ages, there are low levels of platelet activation and the risk genotype may be silent. As platelet activation increases with age, the genotype expresses itself, which leads to stronger age-phenotype associations in the presence of the risk genotype. rs1354034 is a putative functional variant and has been studied before in connection with the platelet-related phenotypes. In particular, rs1354034 disrupts a conserved GATA motif and resides in a regulatory element that has previously been shown to affect the transcription of nearby *ARHGEF3*^51^. Additionally, rs1354034 is associated in trans with the expression of von Willebrand factor (VWF) and other key platelet/megakaryocyte genes found at other loci^52^.

**Figure 7:**
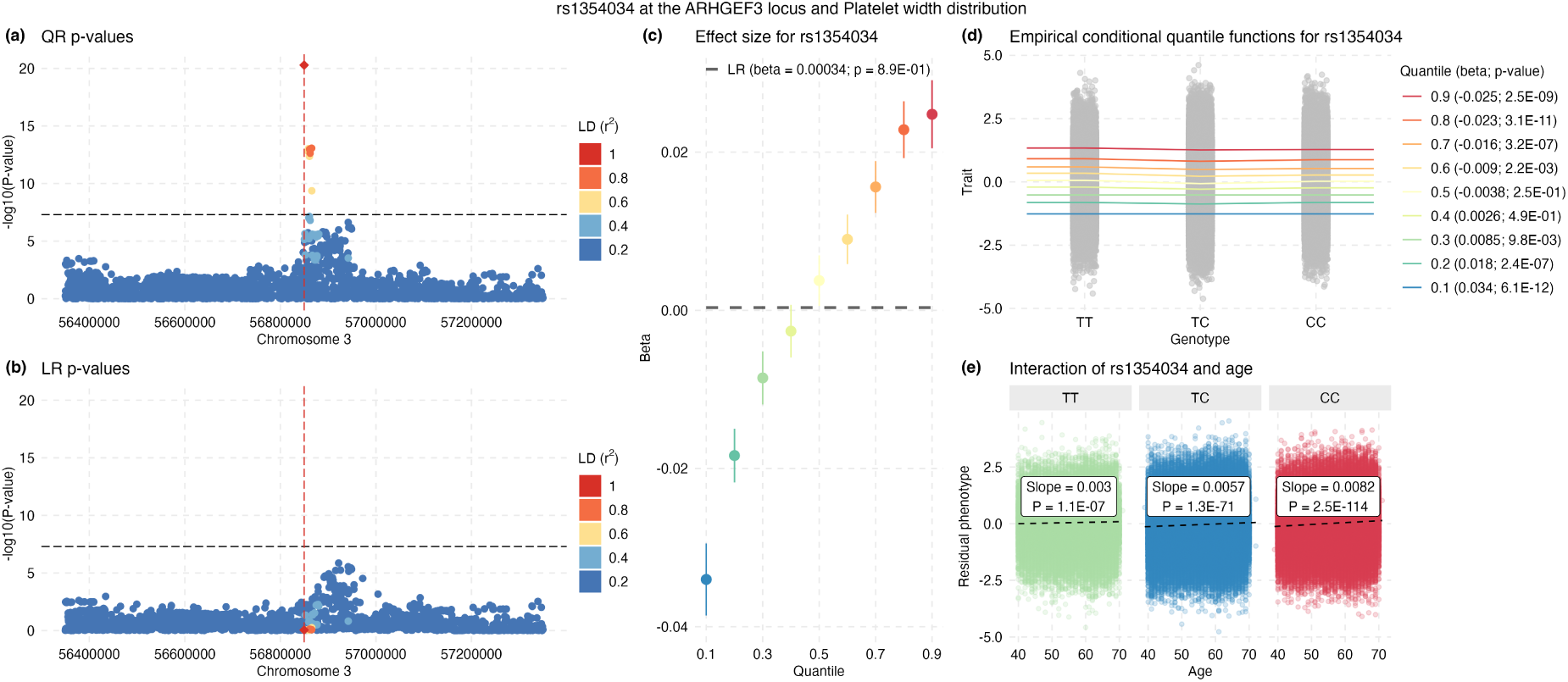
Association of rs1354034 at the *ARHGEF3* locus with Platelet width distribution. (a) -log10(p-value) of QR with the lead QR SNP per locus shown as diamond. (b) -log10(p-value) of LR with the lead QR SNP per locus shown as diamond. Colors indicate the LD (*r*^2^) to the lead SNP for each variant. (c) The estimated quantile specific effects and the SE are shown at *τ* = 0.1, 0.2, 0.3, 0.4, 0.5, 0.6, 0.7, 0.8, 0.9. (d) The curves are the *τ* = 0.1, 0.2, 0.3, 0.4, 0.5, 0.6, 0.7, 0.8, 0.9 empirical conditional quantile functions. Shown in black is also the LR line fitted to the data. (e) gene-environment interaction between rs1354034 and age.

#### Replication of gene-environment interactions

We attempted to replicate the study-wide significant gene-environment interactions above in an independent dataset of European individuals from UKBB (Table S2). Note that the replication datasets are much smaller in size than the discovery datasets. There is strong correlation among estimated effect sizes from the discovery and replication (*r* = 0.77) with almost perfect sign consistency with 56 out of 58 interaction effects showing the same sign (*p* < 2.2*e* 308). In terms of p-value, 25 out of 58 tests are nominally significant (*p* < 2.2*e* 308), with 6 out of 58 being significant after adjusting for 58 tests (*p* < 0.00086), including variants in *TCF7L2* and age for Glycated haemoglobin (HbA1c), variants in *SMOX* and sex for High light scatter reticulocyte count, Reticulocyte percentage, and Immature reticulocyte fraction, variants in *CSP1* and sex for Mean platelet (thrombocyte) volume. Similar results are obtained when relaxing the list of discovered interactions to the ones that are significant for each trait individually (Supplemental Table 1).

## Discussion

Identifying heterogeneous associations as made possible by QR should be of wide interest to genetic association studies of medically relevant traits. We have investigated the potential of QR techniques as a complement to the usual LR for GWAS with biomarkers. QR models can detect distribution-wise associations and efficient implementations in standard statistical packages can readily be applied to GWAS in much the same way as LR. In contrast to LR, QR is robust to non-normality of traits (including heavy-tailed distributions) and outliers, and results from QR are invariant to monotonic trait transformations. QR is more general than the more commonly encountered variance-based tests and can detect effects on the entire phenotype distribution, i.e. beyond variance. Although we have focused on GWAS with independent subjects, QR can be applied to the correlated data setting by including random effects whose covariance structure depends on the genetic relatedness matrix^53, 54^.

In applications to 39 quantitative traits in UKBB we demonstrated the value of performing QR for biomarkers of interest to clinical phenotypes. Overall, the QR approach provides a more complete picture of the distributional effects even for associations detected by LR. Indeed, results from LR are only informative about the effect on the phenotypic mean but cannot tell us what happens in the rest of the conditional distribution. Therefore QR can be used as a complement to LR to help highlight potentially more interesting associations while also providing more detailed information on the associations across different quantile levels. Such knowledge can be very helpful also for functional validation studies as QR associations can point to particular environments where genetic effects are strongest.

SNPs with heterogeneous effects that are detected by QR can arise due to various factors, including gene-gene or gene-environment interactions, time-dependent effects or measurement error. Similarly, genetic nurturing effects^55^, where sequence variants in the parental genomes that are not transmitted to a child interact with the child’s genotype to affect the phenotype, can also lead to heterogeneous effects. Elucidating the precise source of heterogeneity is difficult in general but may be possible by, for example, post-hoc incorporation of explicit interactions (when available) in the model as we have done here, and can suggest particular contexts in which GWAS-discovered variants manifest their effects. For example, interactions with age can suggest that specific genetic associations vary with age, and provide a more nuanced understanding of when in the life-course genetic variants have the strongest effects. Similarly, most of the interactions we detect are with sex, showing that the effect of genetic variants on traits may depend on sex and supporting the importance of considering sex and gender explicitly in genetic association studies.

QR is computationally more expensive than LR, but can be easily scaled up to biobank sized datasets as illustrated here. To give a specific example, for BMI (with 325,306 individuals and 8,572,925 variants), LR with PLINK2^56^ took 92 hours of CPU time and QR (nine quantile-specific tests) with the QRank package in R took 9,534 hours of CPU time. However, both LR and QR can be accelerated by performing tests in parallel on small genomic regions. In our implementation, the genome-wide variant set was split into 1,000 segments of ~ 8.57K variants, with each segment taking about 0.1 hours for LR and 9.5 hours for QR with a single CPU core. With this data scatter and parallel strategy, the elapsed wall-clock time with 1,000 CPU cores became reasonable (∼ 9.5 hours).

Although our focus has been on QR as a natural extension of commonly used LR models for quantitative traits or biomarkers in GWAS, more complex models will be needed to understand the underlying dynamic mechanisms in which genetic effects on a target phenotype are mediated or moderated by multiple other phenotypes that are causally linked with the target. However, a barrier to the development of such models has been the focus of GWAS on single diseases. The advent of large-scale biobank data linked to electronic medical records such as the UKBB make it possible to surmount this barrier by providing a unique opportunity to understand genotype-phenotype associations as a function of age and by accounting for cross-phenotype dependencies. Furthermore, moving beyond marginal models as commonly used is also of interest, and models that take all genetic variants into account are more helpful in allowing us to get to a more causal interpretation of genetic association findings.

In summary, we described QR as a useful complement to LR that can lead to interesting biological discoveries for biomarker traits. QR bears many similarities to LR and can be run efficiently using readily available R packages at biobank scale.

## Methods

We provide first some basic details on the linear and quantile regression models as they relate to the GWAS setting.

### Linear Regression (LR)

We assume that we have *n* independent samples from a population, and for the *i*th subject, we denote by *Y_i_* a continuous (quantitative) phenotype, by *X_i_* = (*X_i_*_1_, *X_ip_*)′ the genotypes at *p* genetic variants, and by *C_i_* = (*C_i_*_1_, *C_iq_*)′ covariates we want to adjust for, such as age, gender and principal components of genetic variation. Furthermore, we denote by *Y* = (*Y*_1_, · · · , *Y_n_*)′ the *n* × 1 sample phenotype vector, by *X* the *n* × *p* genotype matrix, and by *C* the *n* × *q* matrix for covariates. We assume that *Y* is normalized.

First, we consider fitting all *p* variants in a genetic region using a joint linear model:

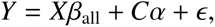

where α ∈ *R^q^*and *β*_all_ ∈ *R^p^* are coefficients, and *ϵ N*(0, σ*_Y_*^2^). We define the error projection matrix *P_C_* = *I C*(*C*′*C*)^−1^*C*′. We project *Y* and *X* onto the orthogonal complement of the column space of *C*, then the estimate for *β*_all_ and its variance can be obtained as

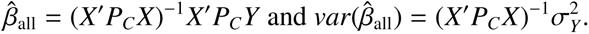

The *z* score test statistic is defined as 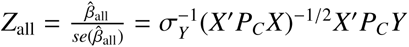.

In GWAS, the typical approach is to perform marginal (unconditional) testing for each variant *j* at a time. Therefore the model becomes:

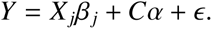

Similarly as above we have:

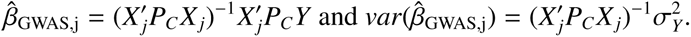

We denote by *β̂*_GWAS_ = (*β̂*_GWAS,1_, · · · , *β̂*_GWAS,p_)′ and the vector of *z* scores as

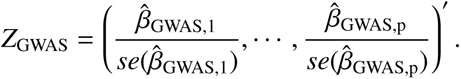

Then, the following is true:

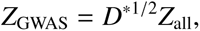

where *D*^∗^ is the correlation matrix of *X*^∗^ := *P_C_ X*. When *X* is independent of *C*, *X*^∗^ = *X* and *D*^∗^ is simply the correlation (LD) matrix among *p* SNPs. Under the null hypothesis *H*_0_ : *β*_all_ = 0, *Z*_all_ ∼ *N*(0, *I_p_*) whereas *Z*_GWAS_ ∼ *N*(0, *D*^∗^).

### Quantile Regression (QR)

We denote the *τ*th conditional quantile function of *Y_i_* as *Q_Yi_* (*τ X_i_*, *C_i_*). Then in linear quantile regression we can write the conditional quantile regression model for *p* joint variants and given a specific quantile level *τ* ∈ (0, 1) as:

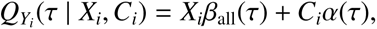

where *β*_all_(*τ*), and α(*τ*) are quantile-specific coefficients and can be estimated by solving the following optimization problem^11^:

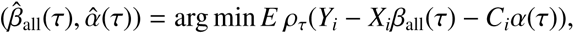

where ρ_*τ*_(*u*) = *u*(*τ − I*(*u* < 0)) is the pinball loss function and *I*(·) is the indicator function. When the error *ϵ* is independent and identically distributed (iid), we have the asymptotic normality for *β̂*_all_(*τ*)^30^ such that

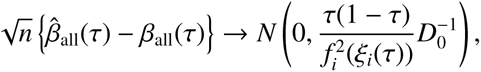

where 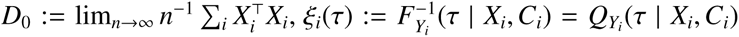, and *f* (·) represents the density function. Thus, the asymptotic distribution of *β^^^_all_* depends on the behavior of the unknown conditional density of the response *Y* in a neighborhood of the conditional quantile model.

For quantile regression, a commonly used hypothesis testing tool is the rank score test^30^. The rank score test statistic for a fixed quantile level *τ* is defined as:

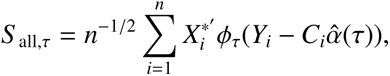

where φ_*τ*_(*u*) = *τ* − *I*(*u* < 0). The covariance matrix of *S _n_*_,*τ*_ is

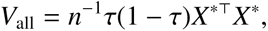

where *X*^∗^ = *P_C_ X* is the same as defined previously. If we fit a marginal model for the *j*th SNP (i.e., QRank^31^), we have

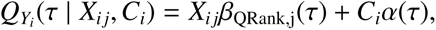

and

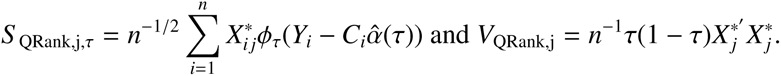

Under the null hypothesis 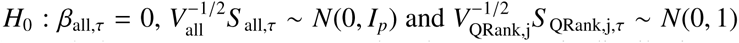, where *V*_QRank,j_ is the *j*th diagonal element of *V*_all_. We note that the asymptotic distribution of the test statistics is independent of the particular distribution of the phenotype. Hence it can be applied to any phenotype without requiring a pre-transformation to achieve normality.

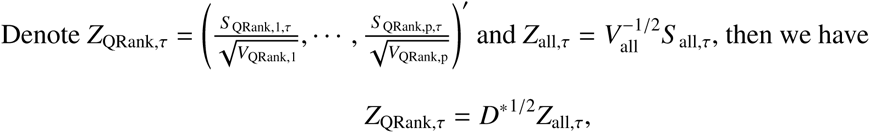

where *D*^∗^ is the correlation matrix of *X*^∗^.

Hence, we conclude that the rank score test statistics from a marginal quantile regression model can be adjusted to the ones from a joint quantile regression model by the correlation matrix *D*^∗^, the same as the *z* scores in linear regression models above. Finally, we combine quantile specific p-values from the marginal tests via Cauchy combination^32^.

### UK Biobank analyses

We used the imputed genotype data of 488,377 participants from the UK Biobank (UKBB). The UKBB samples were directly genotyped with UK BiLEVE and UK Biobank Axiom arrays. Genotype imputation was carried out by using the Haplotype Reference Consortium panel and the merge reference panel from UK10K and 1000 Genomes phase 3 with the IMPUTE4 software^8^. We selected unrelated white British individuals by excluding third-degree or closer relatives (kinship coefficient > 0.0442 estimated with KING^58^) as described in^8, 57^ for discovery analysis and unrelated individuals with non-British European ancestry for replication. The numbers of post-QC samples and variants vary depending on the availability of phenotypic data (Fig. 2). We retained variants with minor allele frequency (MAF) > 1%, Hardy-Weinberg equilibrium (HWE) test p-value > 1 × 10^−6^, and imputation quality (INFO) score > 0.8. Positions of the genetic variants are given in the coordinates of GRCh37 human reference genome.

## Acknowledgements

This study was supported by the National Institutes of Health grants MH095797 (I.I.-L., C.W.) and AG072272 (I.I.-L., Y. W.). The funders had no role in study design, data collection and analysis, decision to publish or preparation of the manuscript. We thank Dan Belsky for useful discussions.

## Competing interests

The authors declare that they have no competing financial interest.

## Data Availability

The manuscript used UK Biobank data available at: https://biobank.ndph.ox.ac.uk/showcase/. The LR and QR GWAS summary statistics for biomarkers in UKBB will be available at Zenodo.

## Code Availability

Scripts for QR GWAS are available at https://github.com/Iuliana-Ionita-Laza/QRGWAS. The quantile regression was performed using the R package quantreg (https://cran.r-project.org/package=quantreg), and the rank score test was performed using the R package QRank (https://cran.r-project.org/package=QRank). The linear regression model was performed using the PLINK 2.0 (https://www.cog-genomics.org/plink/2.0/).

## Supplementary information

**Figure S1:**
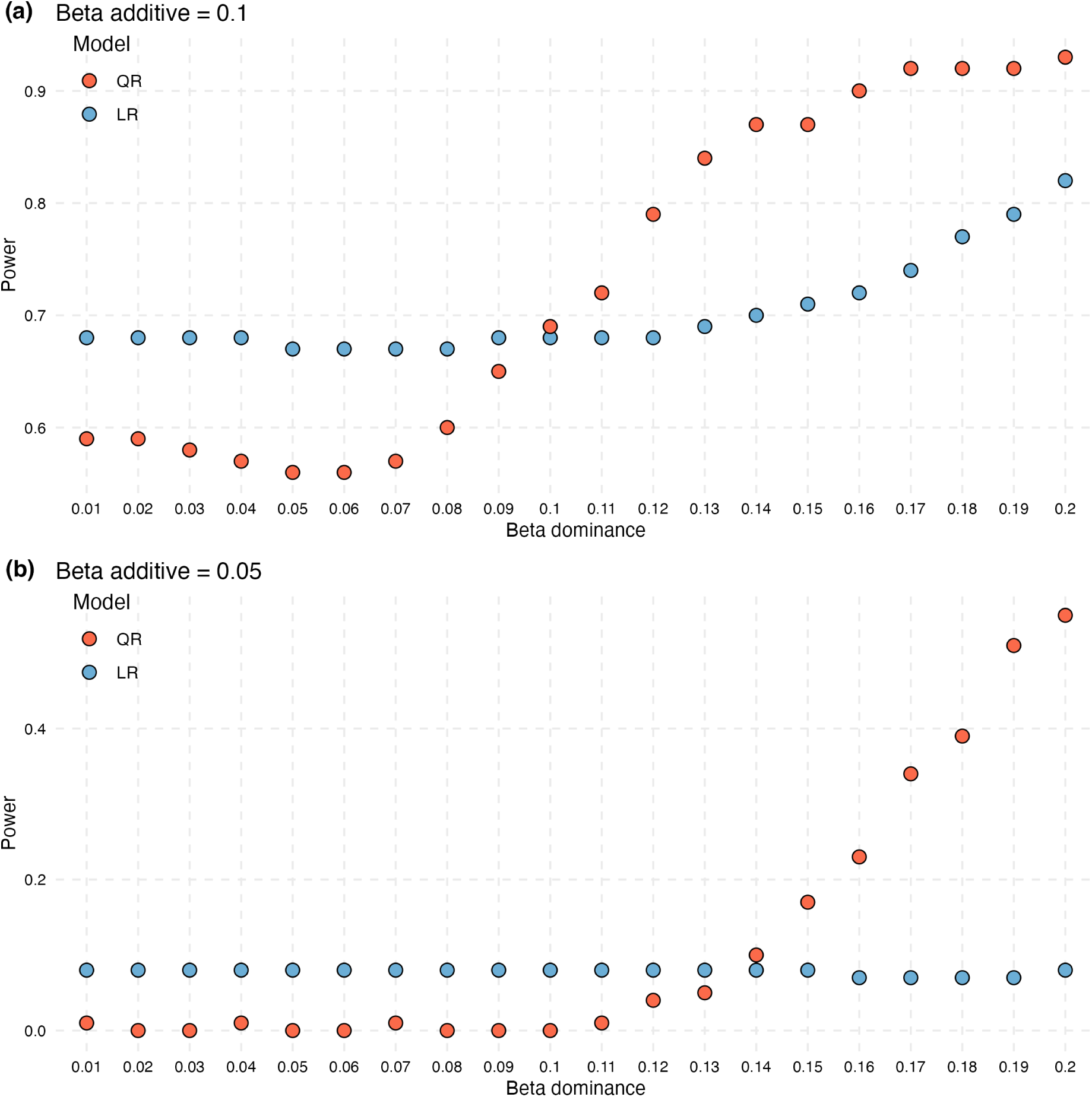
Power for a model with both additive and dominance effects. The additive effect is fixed at 0.1 or 0.05 and the dominance effect varies between 0.01 and 0.2.

**Figure S2:**
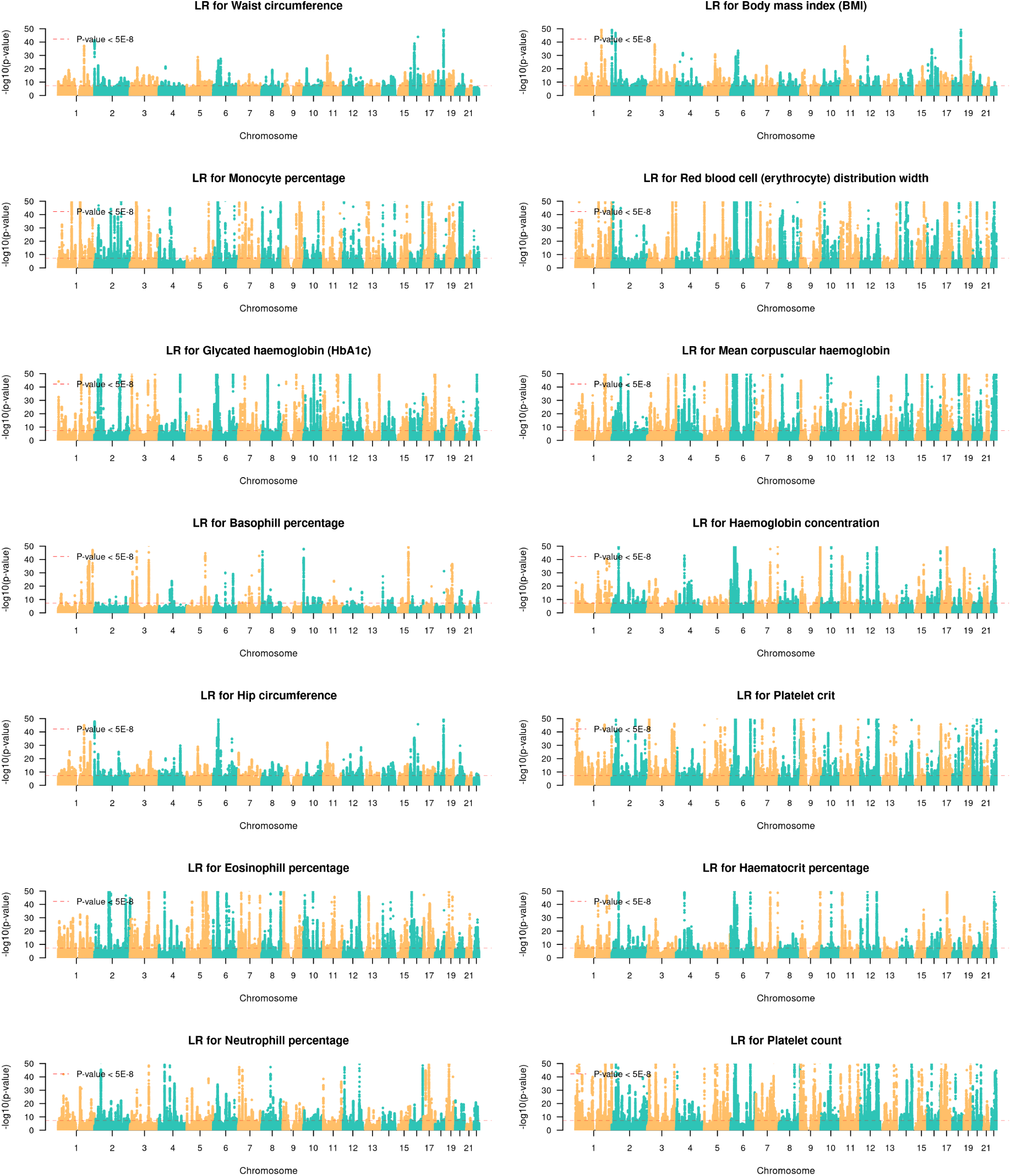
GWAS Manhattan plots for quantitative traits with LR model (Traits 1-14).

**Figure S3:**
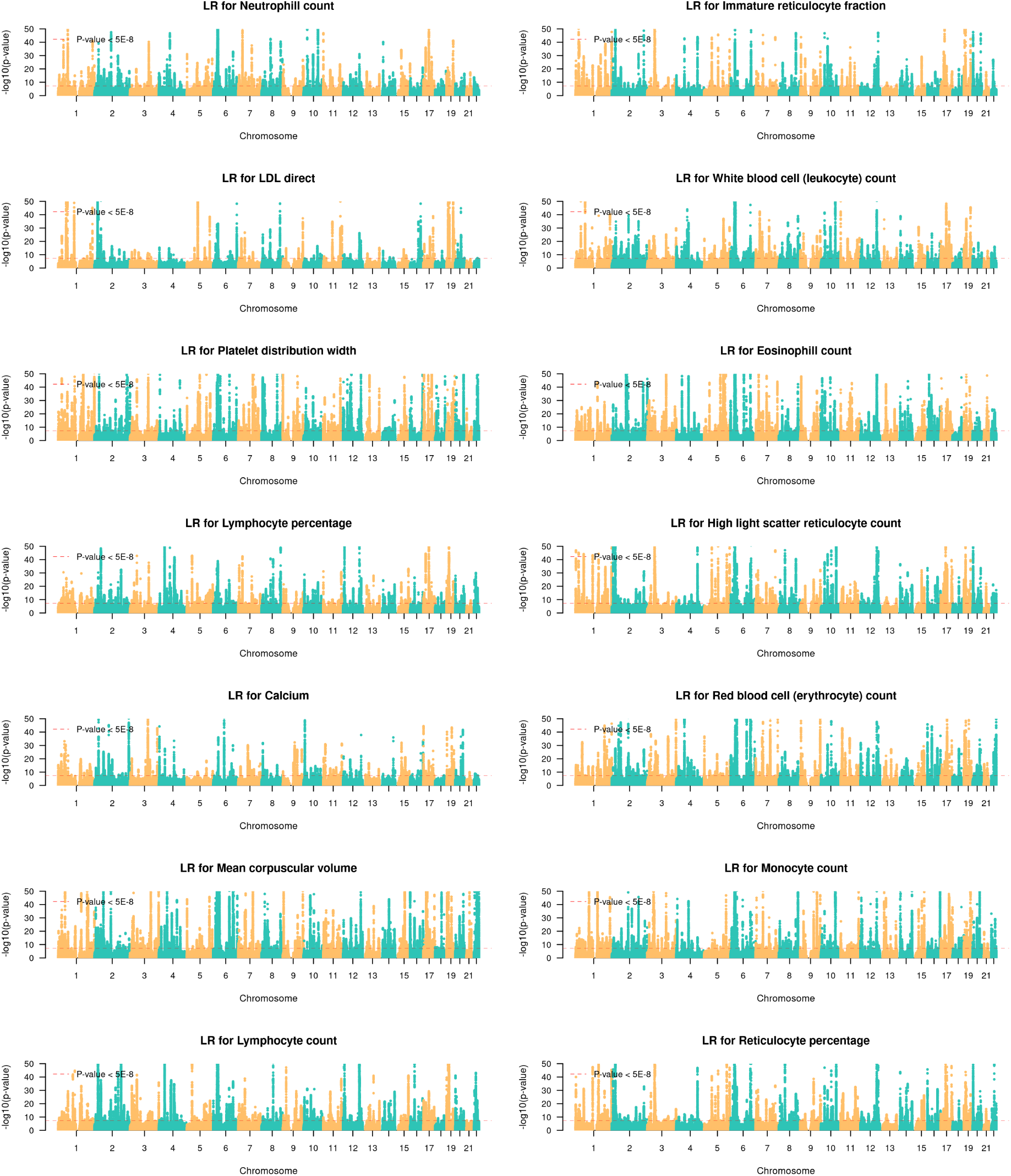
GWAS Manhattan plots for quantitative traits with LR model (Traits 15-28).

**Figure S4:**
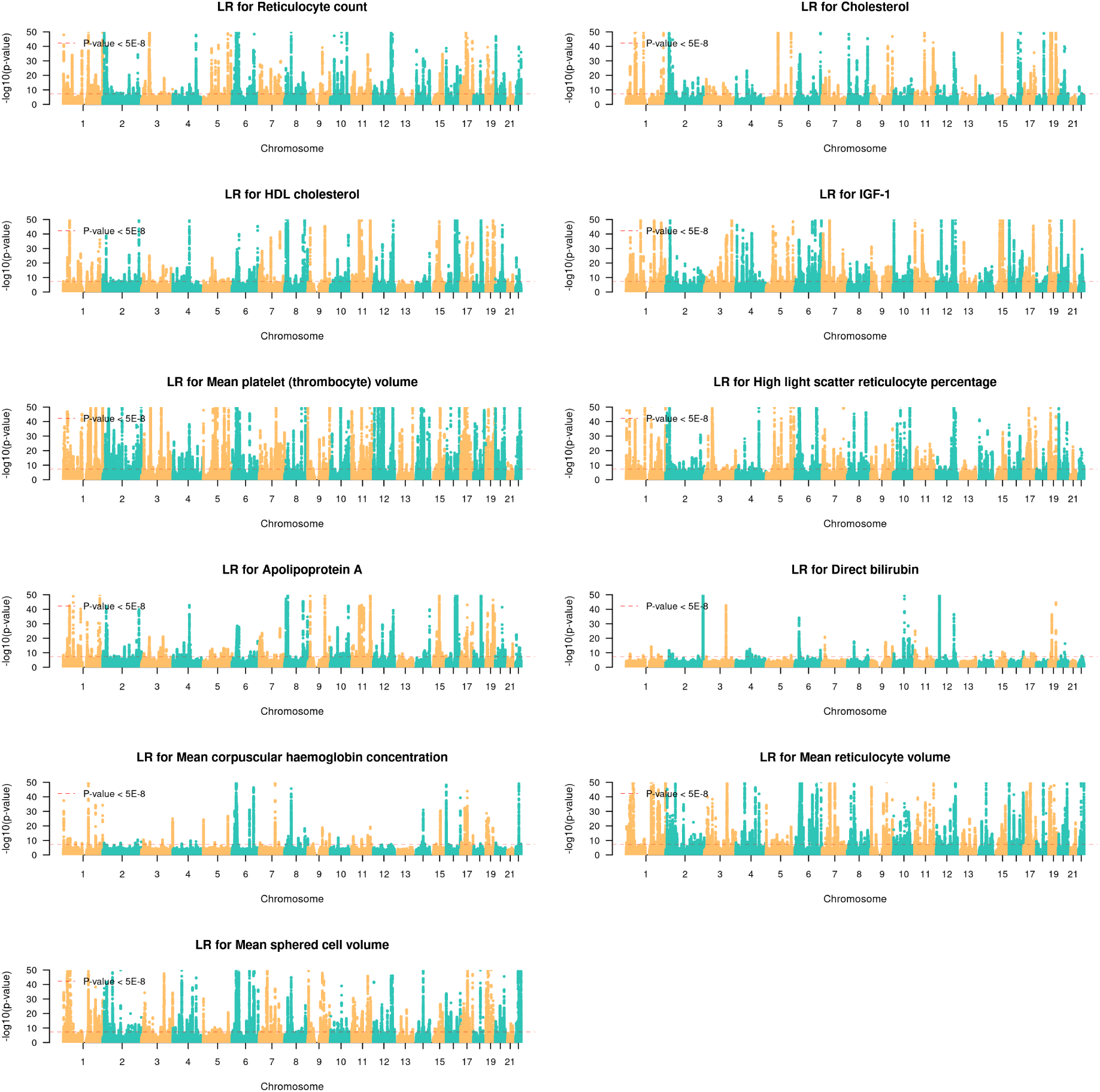
GWAS Manhattan plots for quantitative traits with LR model (Traits 29-39).

**Figure S5:**
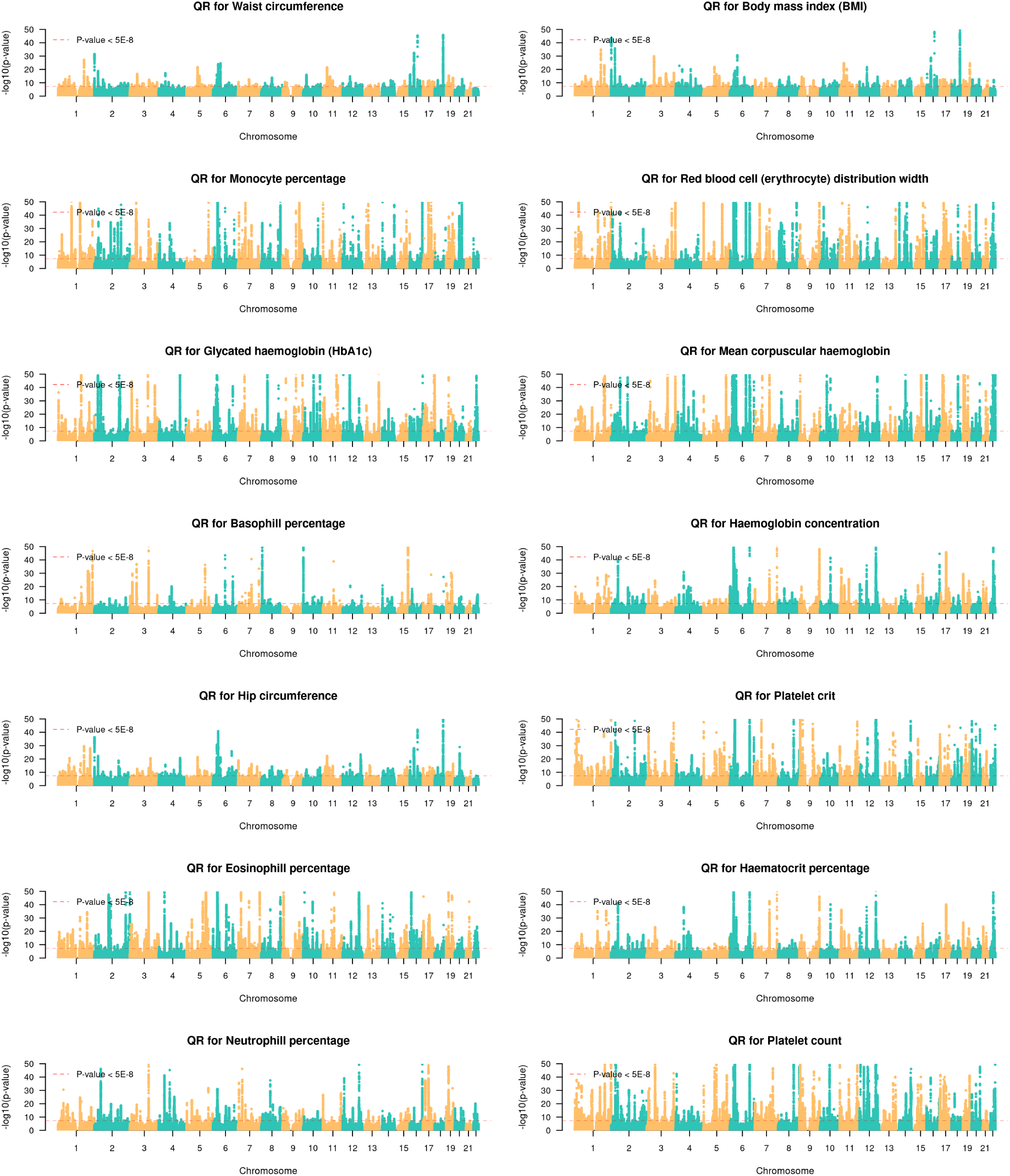
GWAS Manhattan plots for quantitative traits with QR model (Traits 1-14).

**Figure S6:**
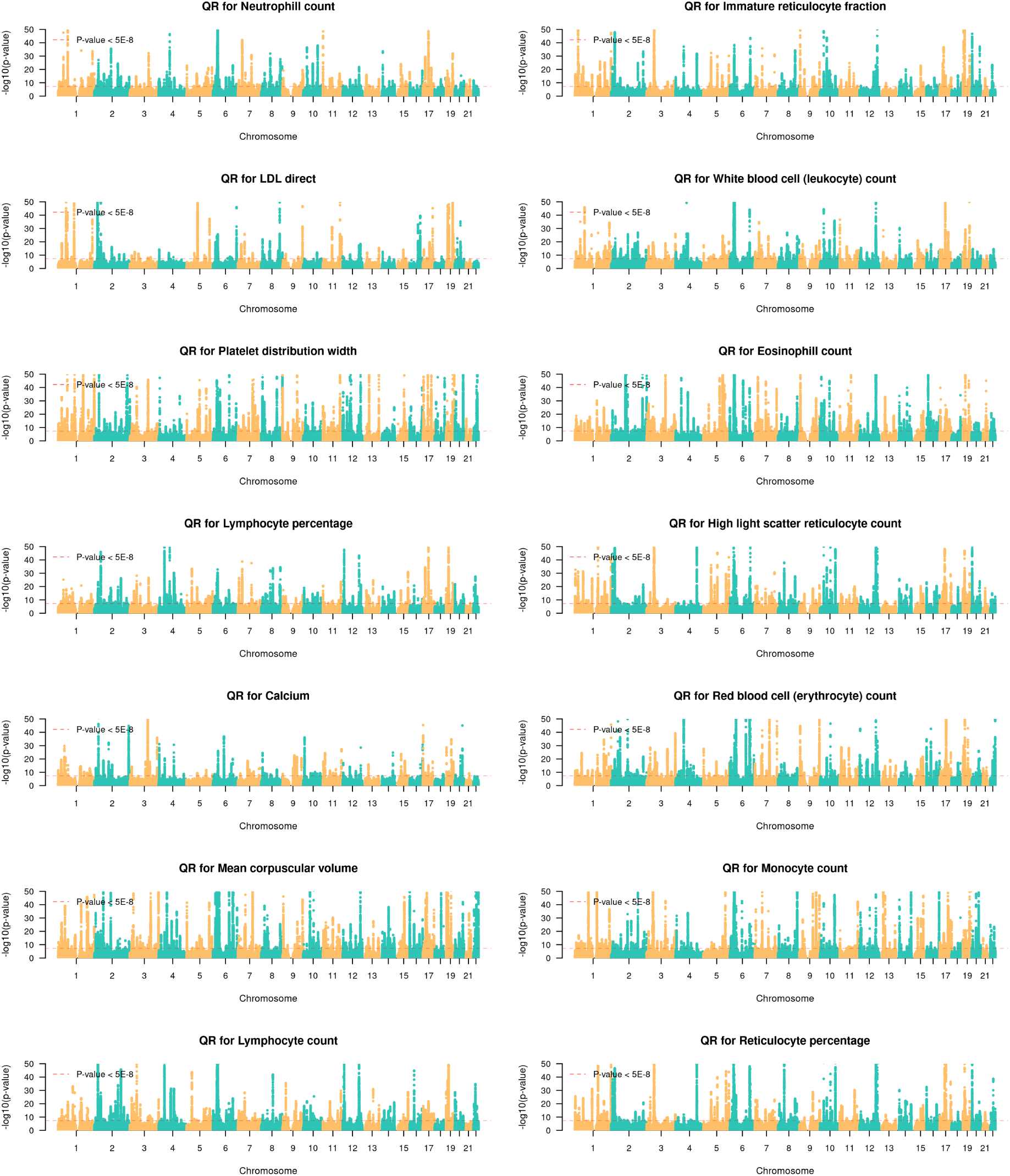
GWAS Manhattan plots for quantitative traits with QR model (Traits 15-28).

**Figure S7:**
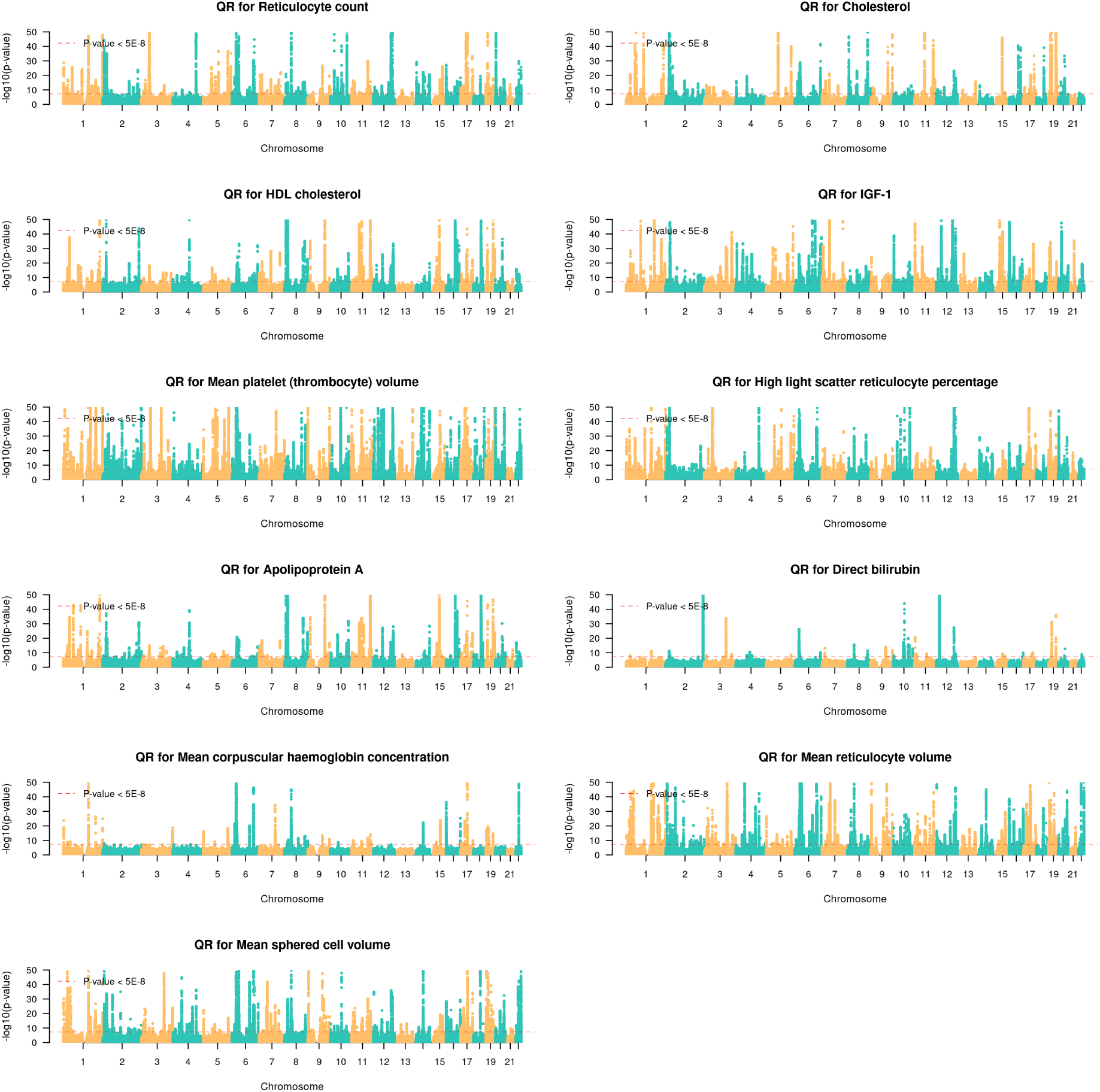
GWAS Manhattan plots for quantitative traits with QR model (Traits 29-39).

**Figure S8:**
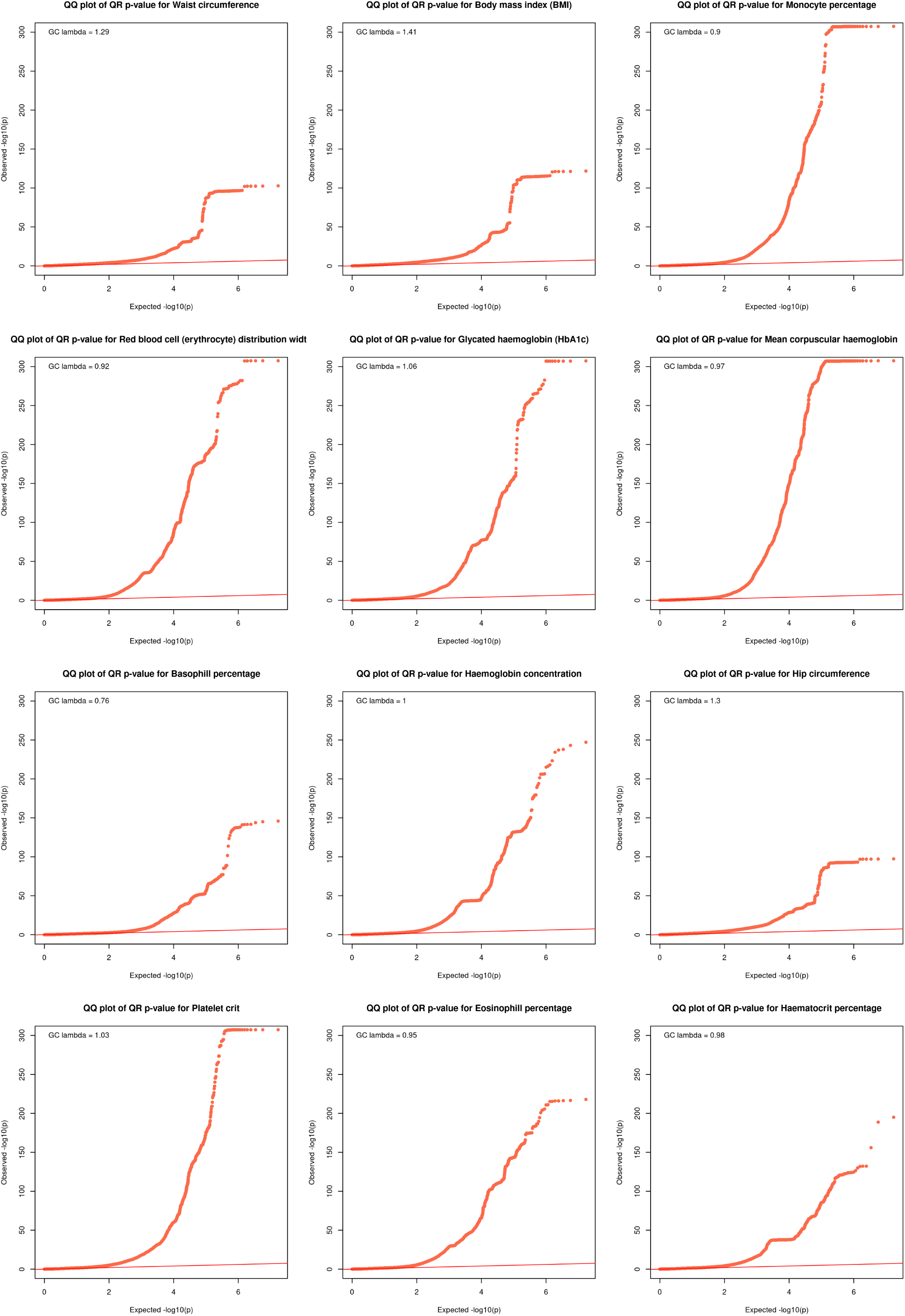
QQ plots of observed versus expected -log10 p-values plots with QR model (Traits 1-12).

**Figure S9:**
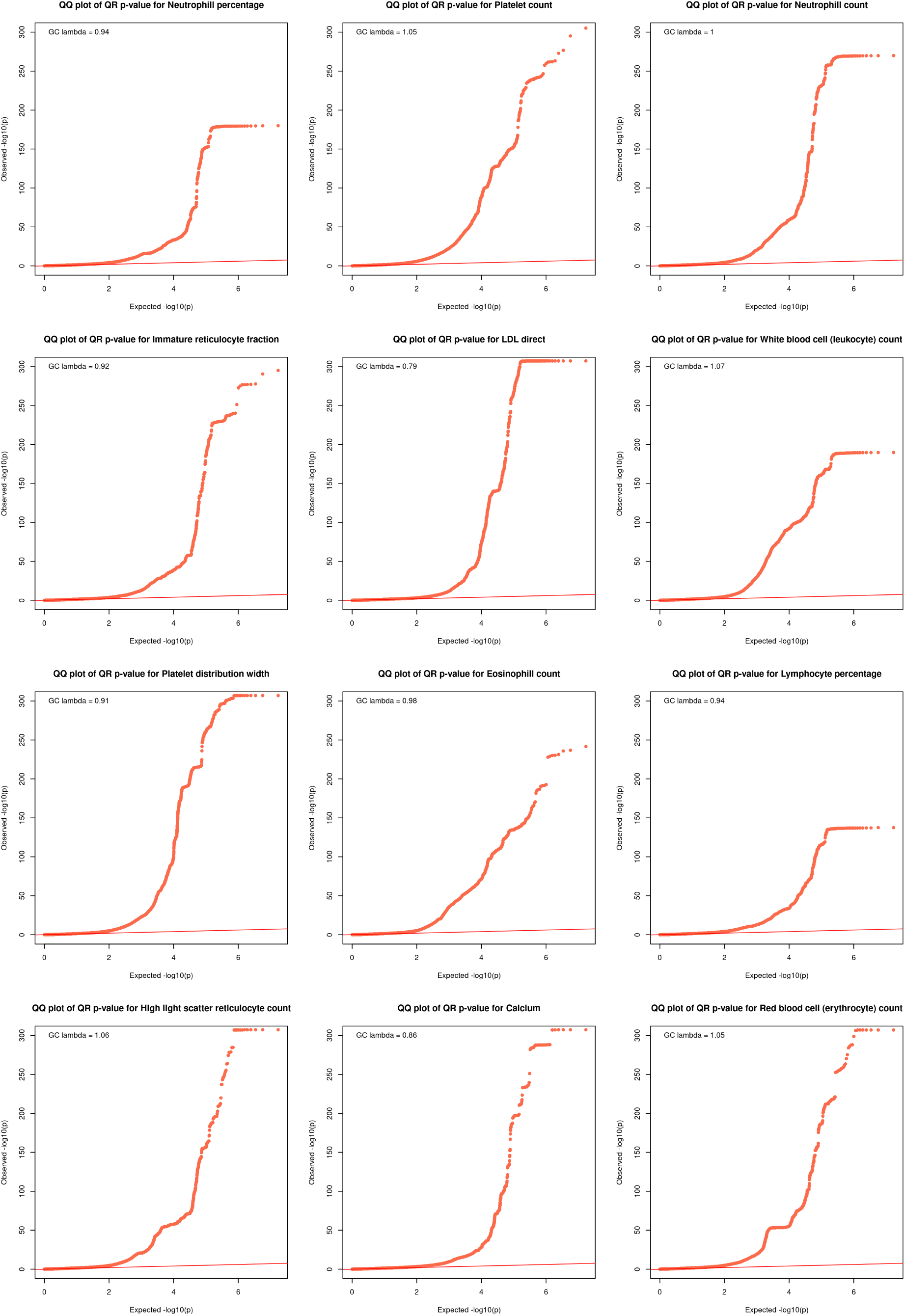
QQ plots of observed versus expected -log10 p-values plots with QR model (Traits 13-24).

**Figure S10:**
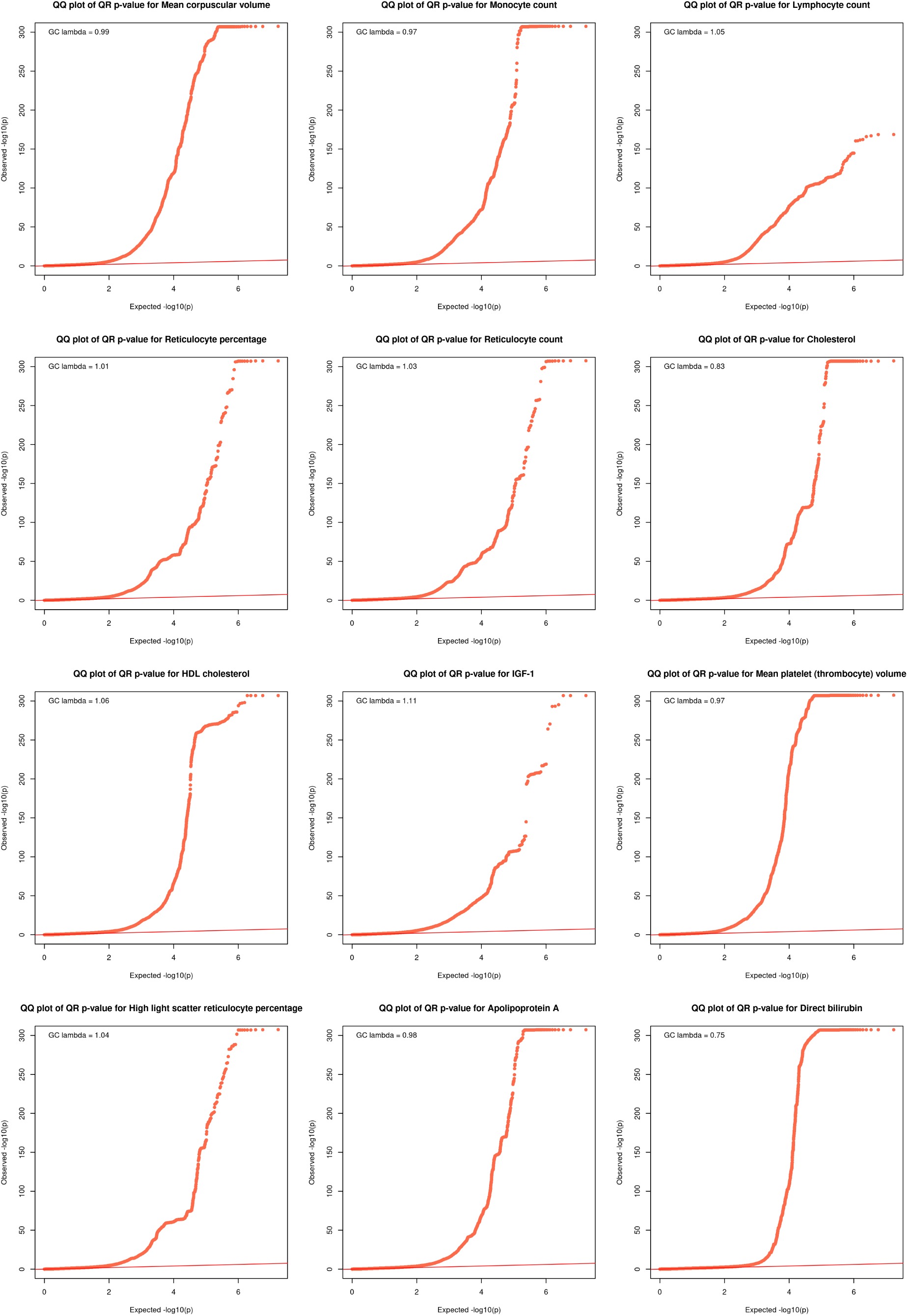
QQ plots of observed versus expected -log10 p-values plots with QR model (Traits 25-36).

**Figure S11:**
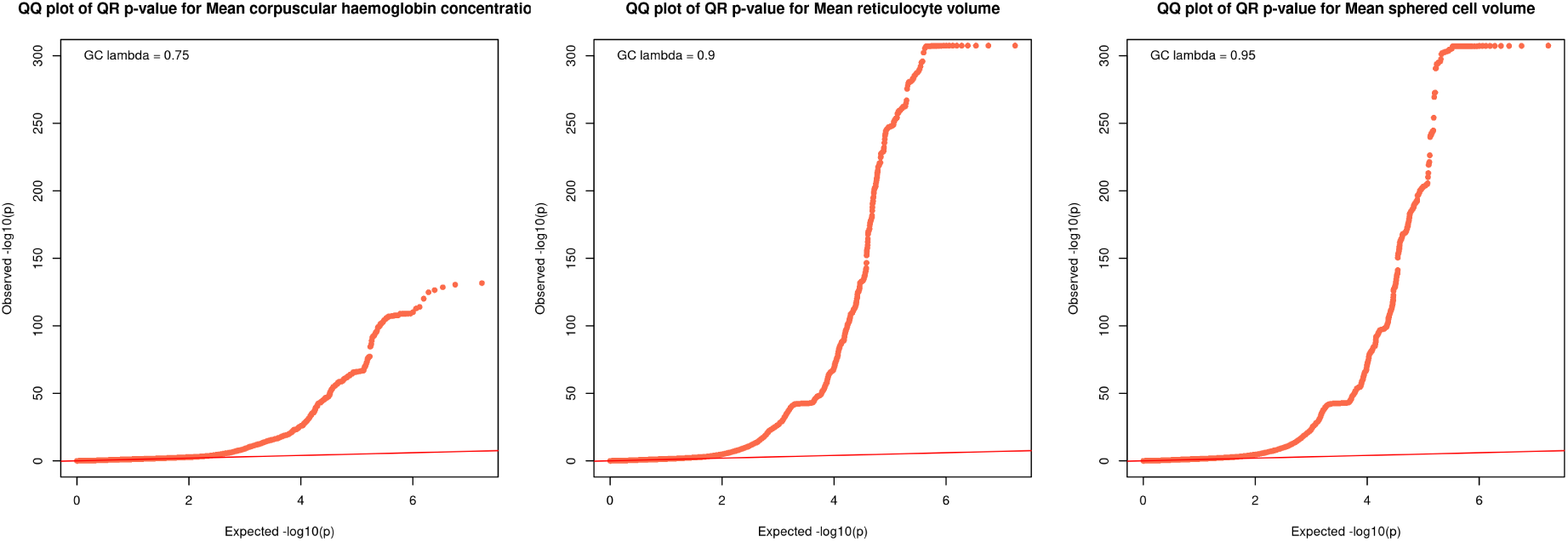
QQ plots of observed versus expected -log10 p-values plots with QR model (Traits 37-39).

**Figure S12:**
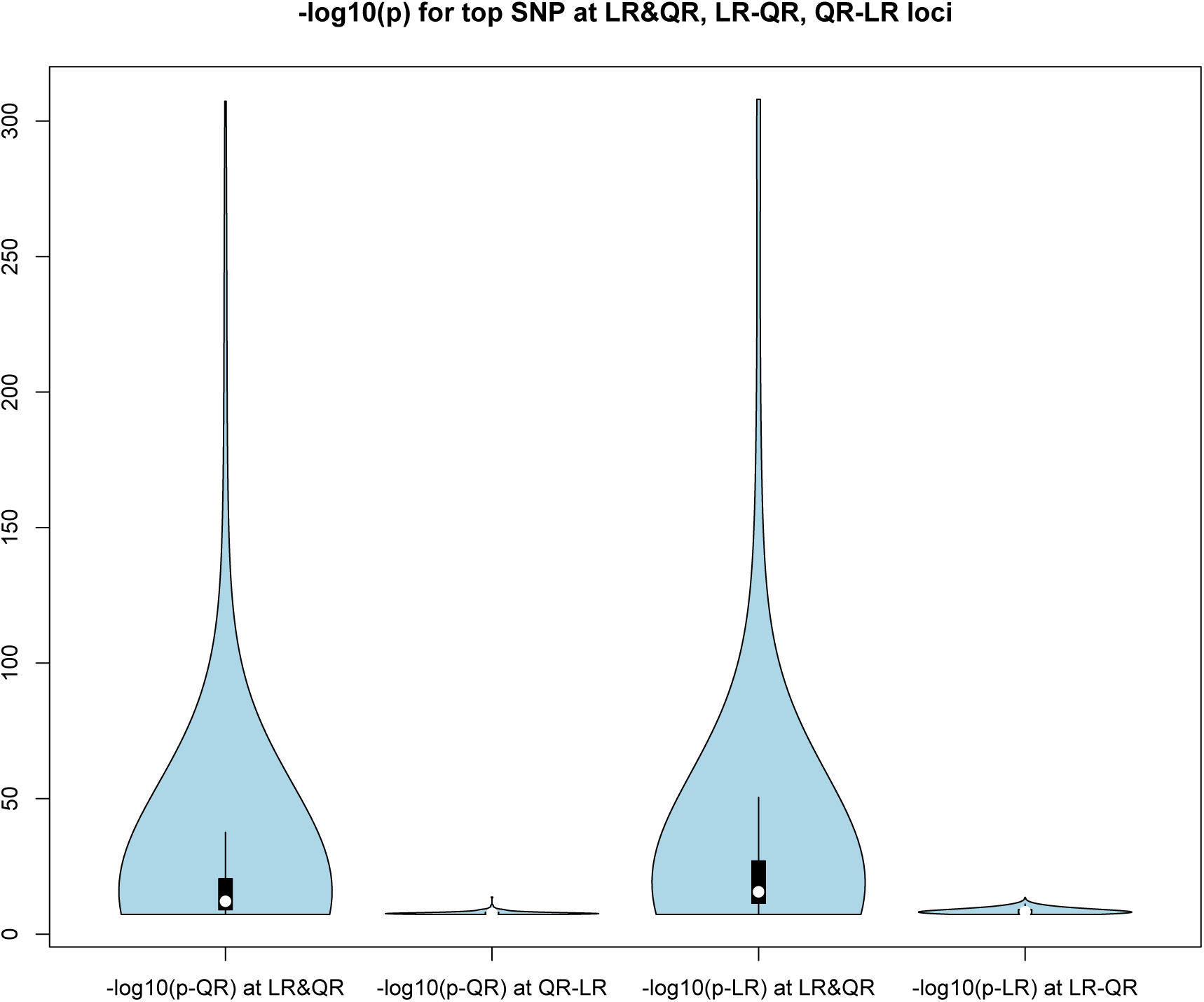
P-value distribution for top SNP at LR&QR, LR-QR and QR-LR loci.

**Figure S13:**
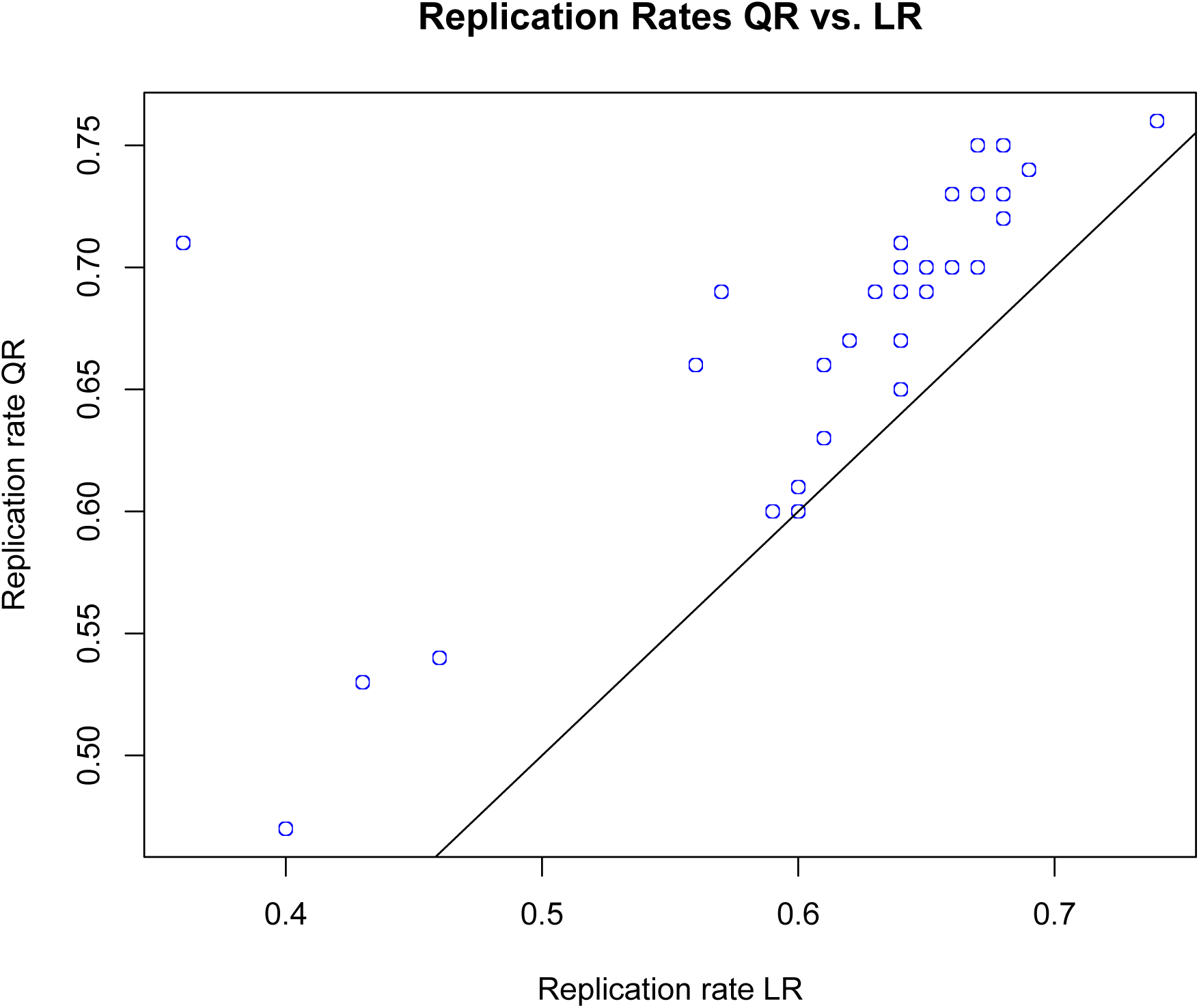
Replication rates for QR vs. LR detected SNPs for 39 traits in UKBB.

**Figure S14:**
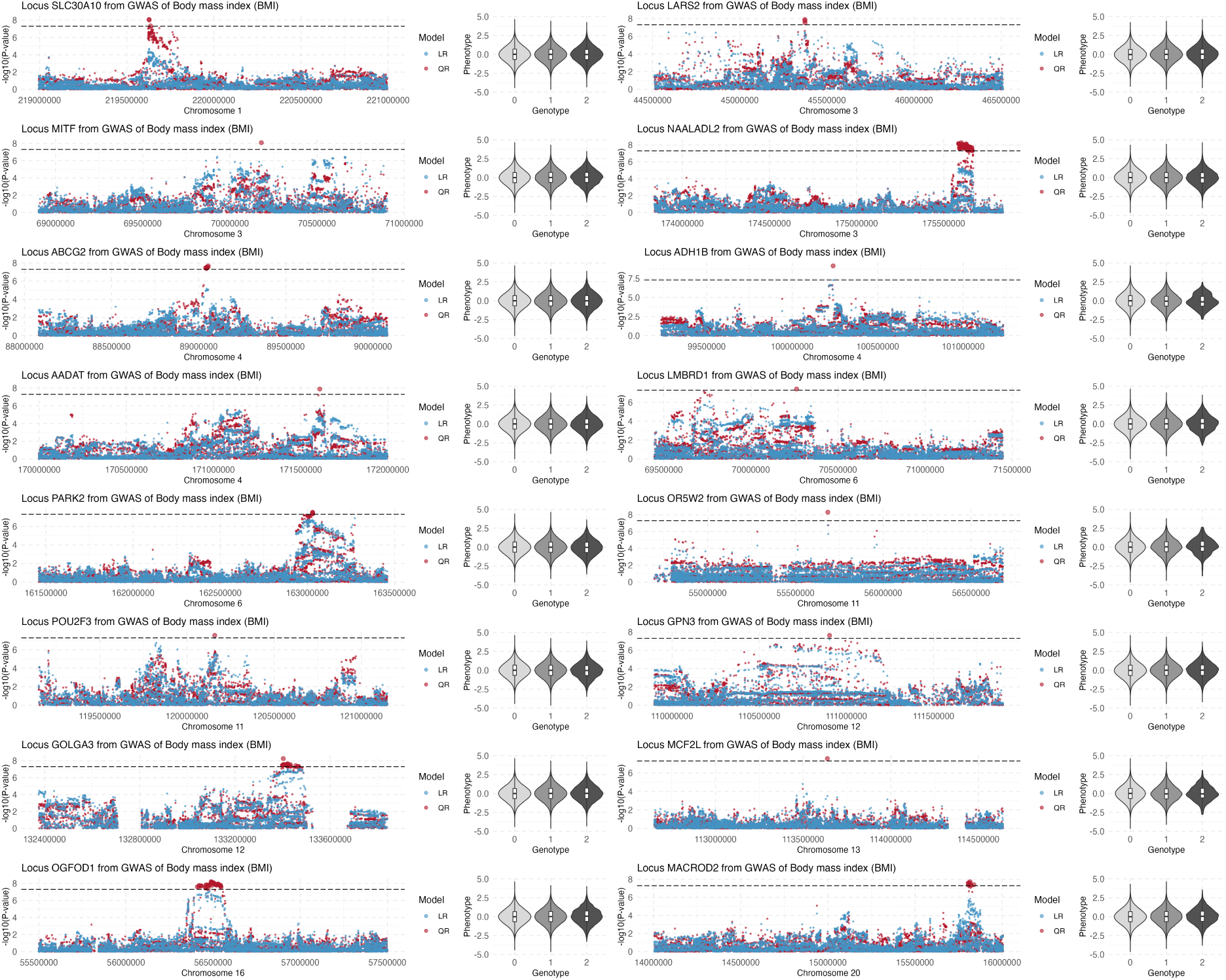
Loci detected by QR-LR for BMI. Manhattan plots with -log10(p-value) for the LR (blue dots) and QR (red dots). For the top QR-LR SNP at each locus, the BMI distributions by genotype group are also shown.

**Figure S15:**
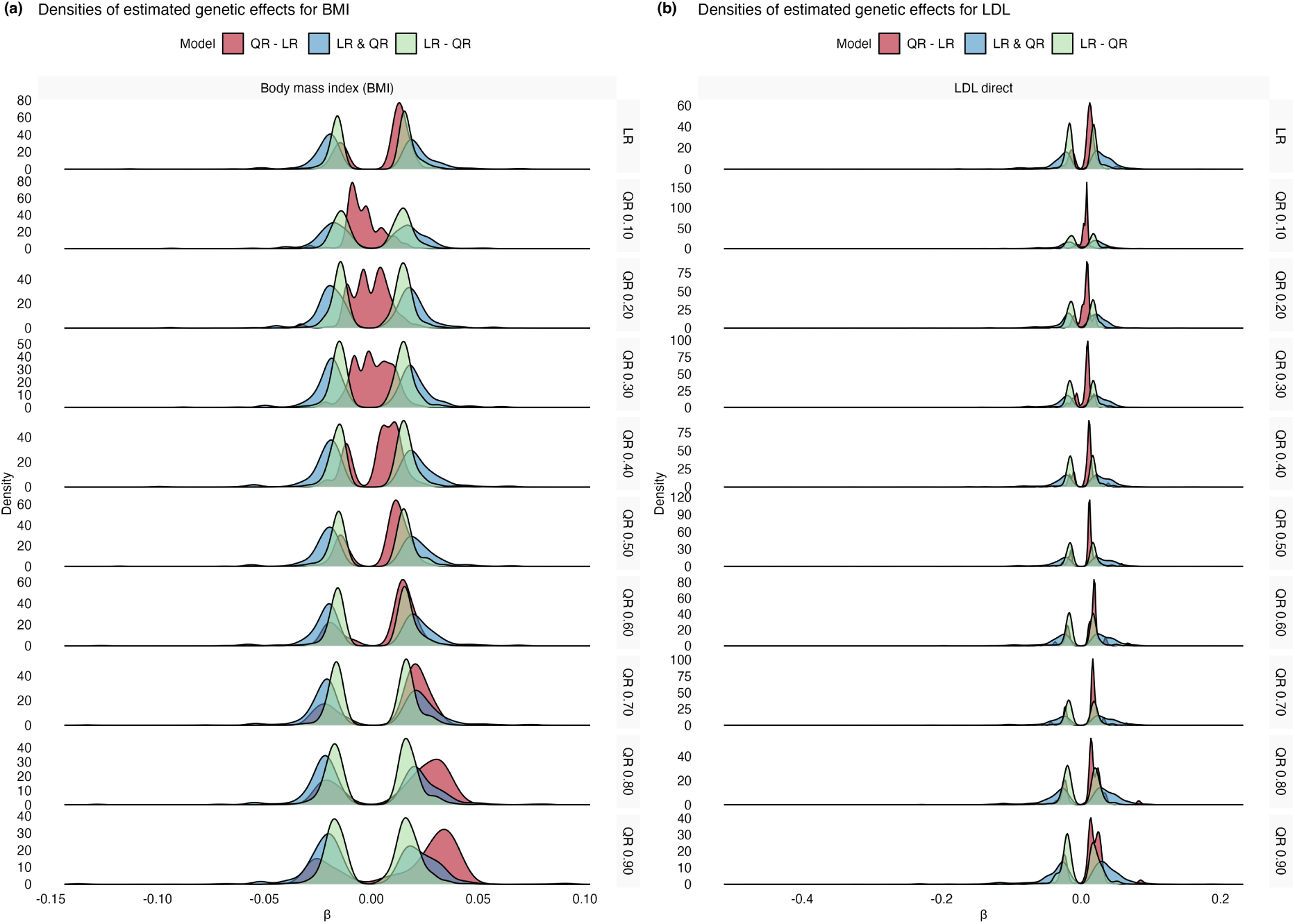
Densities of estimated genetic effects among LR and quantile-specific QR associations for BMI and LDL. Shown in the first row are densities for estimated effects *β*’s from LR for three sets of variants: QR-LR, LR-QR and LR& QR. The quantile-specific rows shows estimated quantile specific coefficients for the same set of SNPs. (a) BMI; (b) LDL.

**Figure S16:**
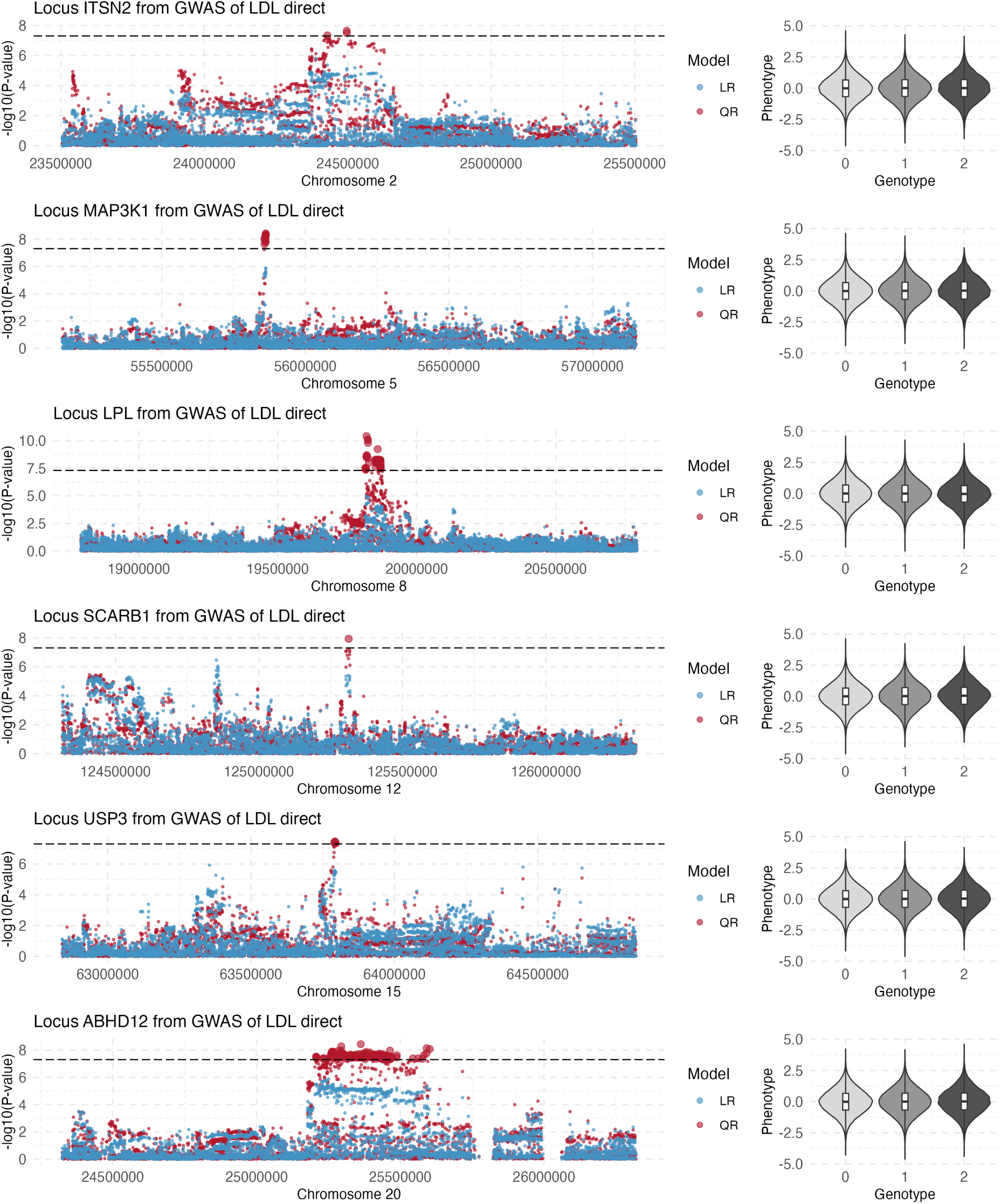
Loci detected by QR-LR for LDL. Manhattan plots with -log10(p-value) for the LR (blue dots) and QR (red dots). For the top QR-LR SNP at each locus, the LDL distributions by genotype group are also shown.

**Figure S17:**
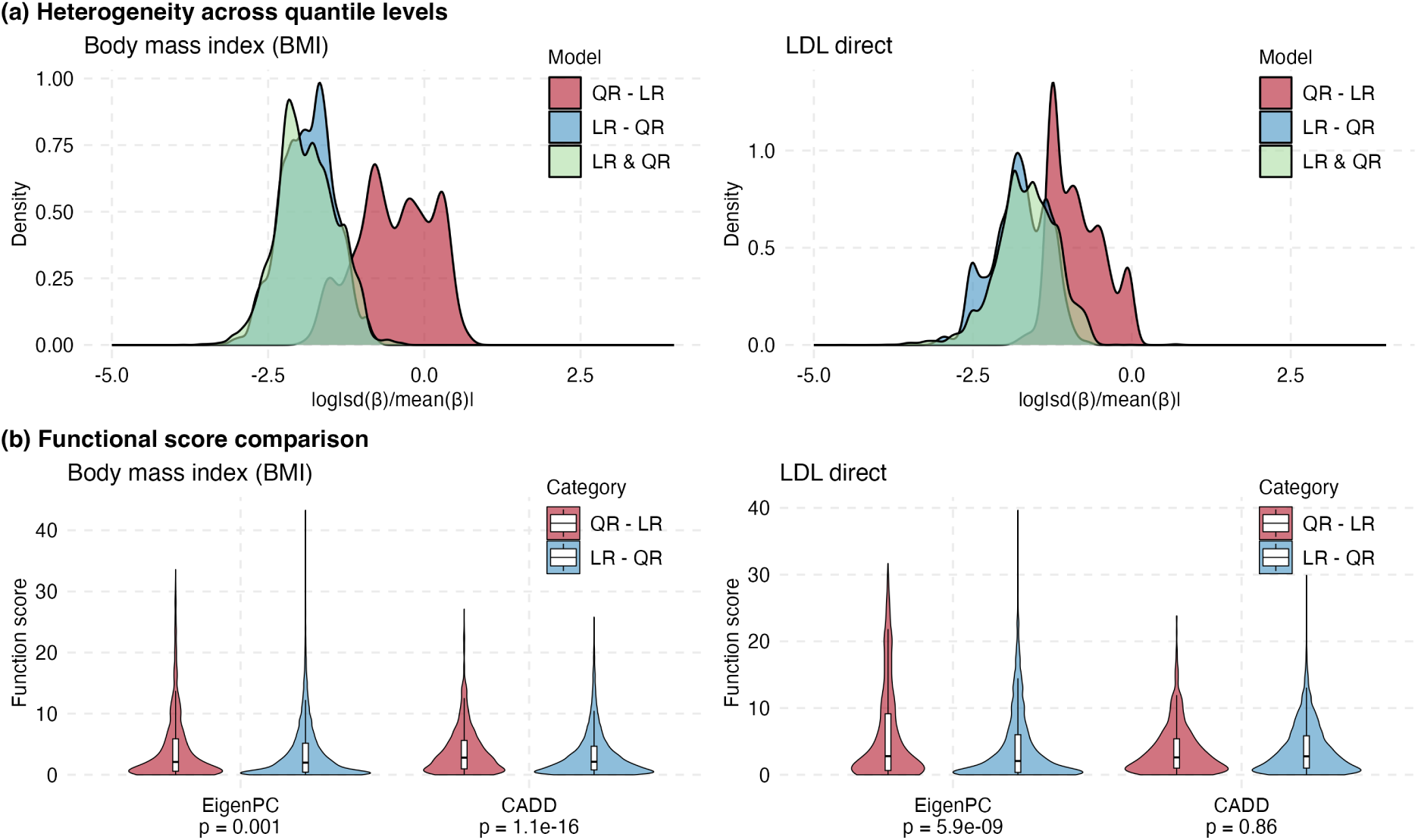
QR-LR vs. LR-QR associations in BMI and LDL. (a) Densities of heterogeneity indexes for GWAS associations in BMI and LDL. Densities are shown for three sets of associated SNPs: LR&QR, LR-QR and QR-LR. (b) Functional scores (EigenPC and CADD) for the associated SNPs detected by QR-LR compared to those from LR - QR. One-sided p-values (QR-LR>LR-QR) from the Kolmogorov-Smirnov test are also shown comparing the functional effects for variants identified by QR-LR vs. LR-QR.

**Figure S18:**
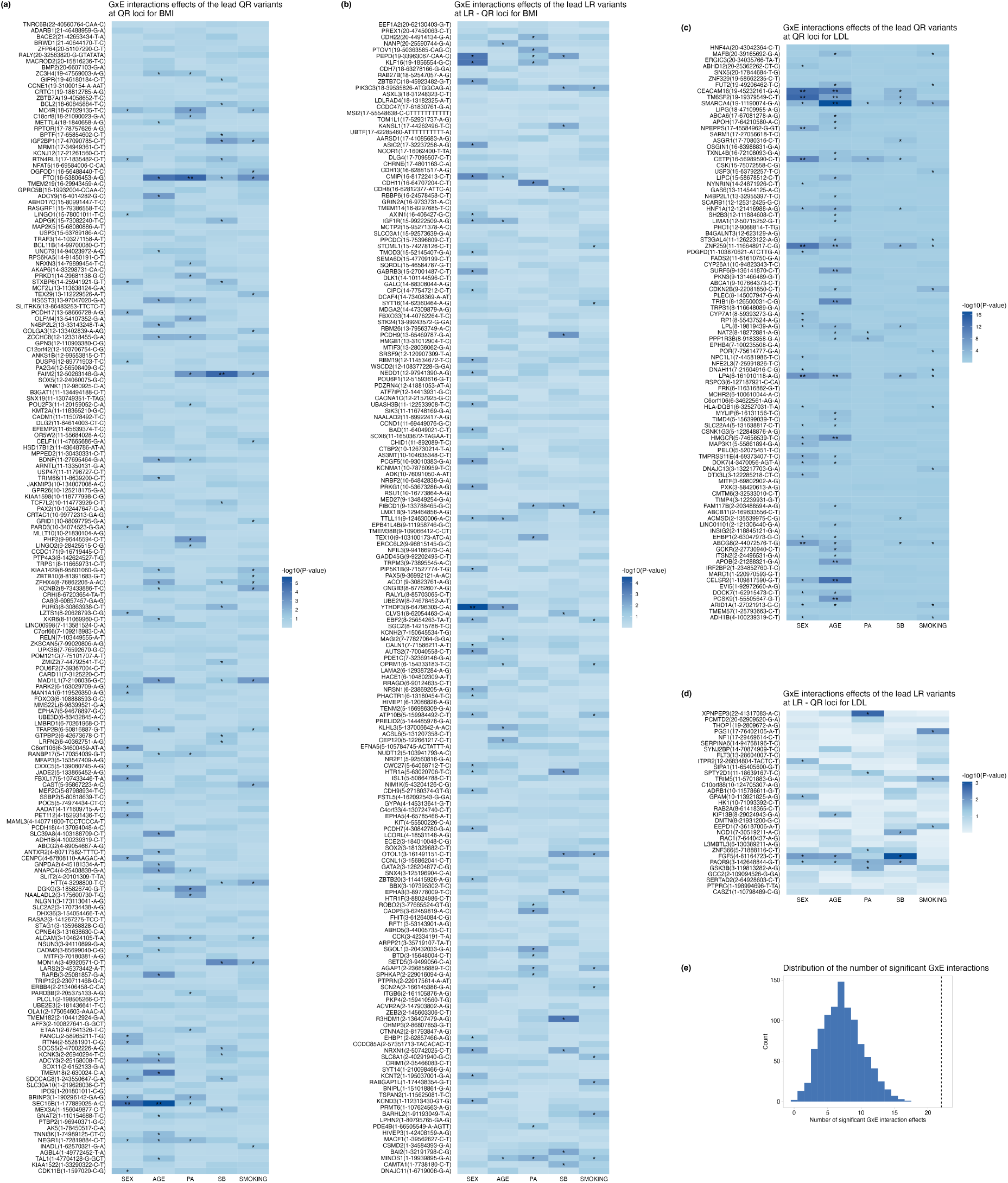
Gene-by-environment interaction effects of the lead QR variants for BMI and LDL. The heatmaps (a-d) show -log10(p-value) of gene-environment interaction tests for the lead QR association at the loci detected by QR, and the lead LR association at loci detected by LR-QR. ”*” denotes nominally significant GxE interaction effects (p-value < 0.05) and ”**” denotes significant effects after Bonferroni correction (*p* < 0.05/(5 281) = 3.55*e* 05). (e) The empirical distribution of the number of significant gene-environment interactions when variants are chosen randomly among significant variants identified by LR or QR.

**Table S1:** Replication rates of QR and LR associations for 39 traits in UKBB. The sample sizes for the replication dataset are also reported. Replications are considered at the nominal level (p-value < 0.05).

| Trait | n | # LR | # QR | % LR Replicated | % LR SignConsistent | % QR Replicated |
| --- | --- | --- | --- | --- | --- | --- |
| Apolipoprotein A | 34542 | 38197 | 26355 | 66.1 | 94.6 | 72.5 |
| Basophil percentage | 38519 | 9502 | 8144 | 56.2 | 95.8 | 66.1 |
| Body mass index (BMI) | 39643 | 38769 | 21357 | 40.4 | 91.5 | 47.2 |
| Calcium | 34768 | 20407 | 13099 | 66.0 | 96.7 | 69.8 |
| Cholesterol | 37961 | 30882 | 24279 | 64.4 | 95.2 | 65.4 |
| Direct bilirubin | 32397 | 11341 | 8819 | 35.7 | 95.6 | 71.3 |
| Eosinophil count | 38518 | 72497 | 59233 | 64.9 | 96.8 | 69.8 |
| Eosinophil percentage | 38519 | 71446 | 59414 | 63.3 | 97.3 | 68.6 |
| Glycated haemoglobin (HbA1c) | 37949 | 62975 | 52460 | 64.9 | 97.3 | 69.0 |
| Haematocrit percentage | 38594 | 47231 | 33668 | 61.2 | 96.3 | 66.5 |
| Haemoglobin concentration | 38594 | 54321 | 44279 | 65.8 | 96.2 | 72.8 |
| HDL cholesterol | 34753 | 41386 | 28812 | 66.9 | 95.9 | 73.4 |
| High light scatter reticulocyte count | 37846 | 59937 | 43311 | 59.6 | 96.5 | 59.6 |
| High light scatter reticulocyte percentage | 37846 | 59229 | 41749 | 58.6 | 96.6 | 60.3 |
| Hip circumference | 39710 | 34372 | 23770 | 45.9 | 91.2 | 54.3 |
| IGF-1 | 37767 | 71071 | 48062 | 61.9 | 96.5 | 67.2 |
| Immature reticulocyte fraction | 37845 | 34104 | 22233 | 56.8 | 96.1 | 68.7 |
| LDL direct | 37876 | 23637 | 17605 | 64.9 | 95.9 | 69.7 |
| Lymphocyte count | 38518 | 65594 | 51217 | 67.7 | 96.7 | 71.5 |
| Lymphocyte percentage | 38519 | 44178 | 31335 | 64.1 | 97.2 | 67.2 |
| Mean corpuscular haemoglobin | 38594 | 77892 | 63110 | 69.5 | 96.9 | 74.2 |
| Mean corpuscular volume | 38594 | 78481 | 61019 | 66.6 | 96.1 | 69.8 |
| Mean corpuscular haemoglobin concentration | 38594 | 19403 | 12177 | 73.8 | 97.3 | 76.4 |
| Mean platelet (thrombocyte) volume | 38594 | 102726 | 77890 | 67.6 | 97.4 | 74.5 |
| Mean reticulocyte volume | 37845 | 64160 | 50139 | 68.3 | 97.3 | 71.9 |
| Mean spheroid cell volume | 37846 | 58204 | 42187 | 63.9 | 97.1 | 70.2 |
| Monocyte count | 38518 | 61928 | 50652 | 61.2 | 95.9 | 62.9 |
| Monocyte percentage | 38519 | 49586 | 38871 | 67.3 | 95.7 | 75.4 |
| Neutrophil count | 38518 | 53651 | 38577 | 64.2 | 96.2 | 64.7 |
| Neutrophil percentage | 38519 | 38690 | 28856 | 67.2 | 97.2 | 70.0 |
| Platelet count | 38594 | 82682 | 59927 | 64.3 | 97.2 | 69.5 |
| Platelet crit | 38594 | 62486 | 43461 | 64.5 | 96.1 | 71.1 |
| Platelet distribution width | 38594 | 66765 | 48286 | 63.4 | 96.5 | 68.9 |
| Red blood cell (erythrocyte) count | 38594 | 58860 | 41669 | 65.3 | 96.1 | 69.7 |
| Red blood cell (erythrocyte) distribution width | 38594 | 73842 | 60566 | 68.3 | 96.8 | 73.3 |
| Reticulocyte count | 37846 | 57534 | 42411 | 60.8 | 96.1 | 63.2 |
| Reticulocyte percentage | 37846 | 57737 | 41767 | 60.4 | 96.7 | 61.2 |
| Waist circumference | 39714 | 23402 | 12019 | 42.8 | 89.9 | 52.8 |
| White blood cell (leukocyte) count | 38593 | 60413 | 44884 | 64.2 | 95.8 | 69.9 |

**Table S2:**
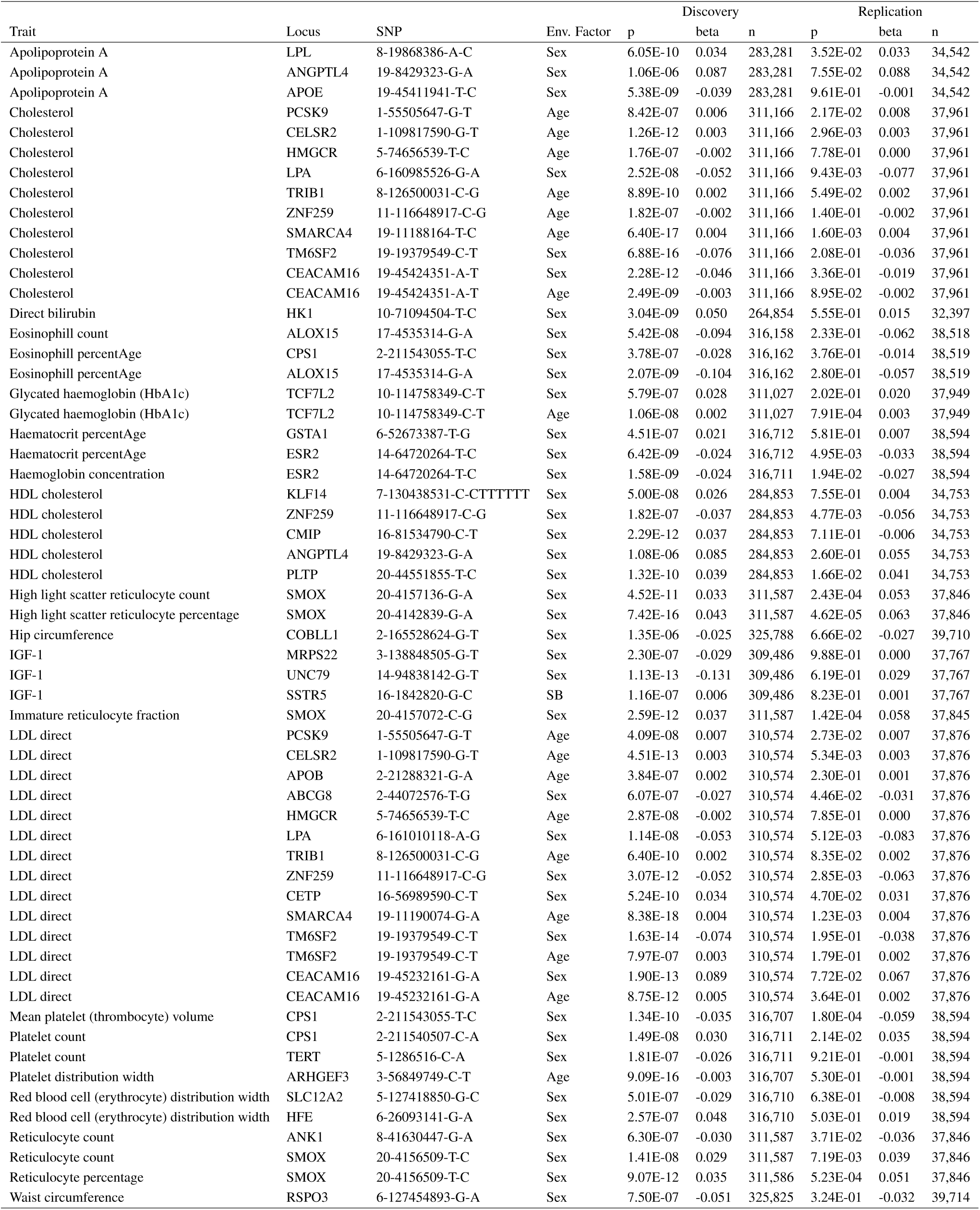
Significant gene-environment interactions using top QR SNP at each locus from QR GWAS for 39 traits. The discovery data includes White British individuals from UKBB, while the independent replication dataset includes European individuals also from UKBB. The sample size *n*, the estimated effect size for the interaction effect *beta* and the p-value for the gene-environment test *p* are shown. Environmental factors include Sex, Age, Sedentary Behavior (SB), Physical Activity (PA) and smoking.

## Notes

### Competing Interest Statement

The authors have declared no competing interest.

## References

1. P. M. Visscher, N. R. Wray, Q. Zhang, P. Sklar, M. I. McCarthy, M. A. Brown, and J. Yang. 10 years of GWAS discovery: biology, function, and translation. The American Journal of Human Genetics, 101: 5–22 (2017).

2. S. Chun, A. Casparino, N. A. Patsopoulos, D. C. Croteau-Chonka, B. A. Raby, P. L. De Jager, S. R. Sunyaev, and C. Cotsapas. Limited statistical evidence for shared genetic effects of eQTLs and autoimmune-disease-associated loci in three major immune-cell types. Nature genetics, 49(4):600–605 (2017).

3. B. D. Umans, A. Battle, and Y. Gilad. Where are the disease-associated eQTLs? Trends in Genetics, 37(2):109–124 (2021).

4. H. Izgi, D. Han, U. Isildak, S. Huang, E. Kocabiyik, P. Khaitovich, M. Somel, and H. M. Dönertacs. Inter-tissue convergence of gene expression during ageing suggests age-related loss of tissue and cellular identity. Elife, 11:e68048 (2022).

5. J. T. Leek and J. D. Storey. Capturing heterogeneity in gene expression studies by surrogate variable analysis. PLoS genetics, 3(9):e161 (2007).

6. M. Somel, P. Khaitovich, S. Bahn, S. Pääbo, and M. Lachmann. Gene expression becomes heterogeneous with age. Current Biology, 16(10):R359–R360 (2006).

7. E. Budinska, V. Popovici, S. Tejpar, G. D’Ario, N. Lapique, K. O. Sikora, A. F. Di Narzo, P. Yan, J. G. Hodgson, S. Weinrich, et al. Gene expression patterns unveil a new level of molecular heterogeneity in colorectal cancer. The Journal of pathology, 231(1):63–76 (2013).

8. Bycroft C, Freeman C, Petkova D et al. The UK Biobank resource with deep phenotyping and genomic data. Nature. 562: 203–209 (2018).

9. Loh, P.-R. et al. Efficient Bayesian mixed model analysis increases association power in large cohorts. Nat Genet. 47, 284–290 (2015).

10. Jiang L, Zheng Z, Qi T, Kemper KE, Wray NR, Visscher PM, Yang J. A resource-efficient tool for mixed model association analysis of large-scale data. Nat Genet. 51: 1749–1755 (2019).

11. Koenker R and Bassett G JR. Regression quantiles. Econometrica. 46: 33–50 (1978).

12. Yang J et al. FTO genotype is associated with phenotypic variability of body mass index. Nature. 490: 267–272 (2012).

13. Brown AA, Buil A, Viñuela A, Lappalainen T, Zheng HF, Richards JB, Small KS, Spector TD, Dermitzakis ET, Durbin R. Genetic interactions affecting human gene expression identified by variance association mapping. eLife 3:e01381 (2014).

14. Pare G, Cook NR, Ridker PM, Chasman DI. On the use of variance per genotype as a tool to identify quantitative trait interaction effects: A report from the Women’s Genome Health Study. PLoS Genet. 6 e1000981 (2010).

15. Song X, Li G, Zhou Z, Wang X, Ionita-Laza I, Wei Y. QRank: A novel quantile regression tool for eQTL discovery. Bioinformatics. 33: 2123–2130 (2017).

16. Wang H, Zhang F, Zeng J, Wu Y, Kemper KE, Xue A, Zhang M, Powell JE, Goddard ME, Wray NR, Visscher PM, McRae AF, Yang J. Genotype-by-environment interactions inferred from genetic effects on phenotypic variability in the UK Biobank. Sci Adv. 2019 Aug 14;5(8):eaaw3538. doi: 10.1126/sciadv.aaw3538.

17. Westerman KE, Majarian TD, Giulianini F, Jang DK, Miao J, Florez JC, Chen H, Chasman DI, Udler MS, Manning AK, Cole JB. Variance-quantitative trait loci enable systematic discovery of gene-environment interactions for cardiometabolic serum biomarkers. Nat Commun.13: 3993 (2022).

18. Manchia M, Cullis J, Turecki G, Rouleau GA, Uher R, Alda M. The impact of phenotypic and genetic heterogeneity on results of genome wide association studies of complex diseases. PLoS ONE 8 e76295 (2013).

19. van Ijzendoorn MH, Bakermans-Kranenburg MJ, Belsky J, Beach S, Brody G, Dodge KA, Greenberg M, Posner M, Scott S. Gene-by-environment experiments: a new approach to finding the missing heritability. Nat Rev Genet. 2011 Nov 18;12(12):881; author reply 881. doi: 10.1038/nrg2764-c1.

20. Palmer DS, Zhou W, Abbott L, Wigdor EM, Baya N, Churchhouse C, Seed C, Poterba T, King D, Kanai M, Bloemendal A, Neale BM. Analysis of genetic dominance in the UK Biobank. Science. 379(6639): 1341-1348 (2023)

21. Findley AS, Monziani A, Richards AL, Rhodes K, Ward MC, Kalita CA, Alazizi A, Pazokitoroudi A, Sankararaman S, Wen X, Lanfear DE, Pique-Regi R, Gilad Y, Luca F. Functional dynamic genetic effects on gene regulation are specific to particular cell types and environmental conditions. Elife. 2021 May 14;10:e67077. doi: 10.7554/eLife.67077.

22. Levene H. Robust tests for equality of variances. Contributions to Probability and Statistics: Essays in Honor of Harold Hotelling (Stanford Univ. Press) 278–292 (1960).

23. Brown MB, Forsythe AB. Robust tests for the equality of variances. J. Am. Stat. Assoc., 69: 364–367 (1974).

24. Brown A.A. et al. Genetic interactions affecting human gene expression identified by variance association mapping. Elife, 3, e01381 (2014).

25. Smyth GK. Generalized linear models with varying dispersion. J. R. Stat. Soc. B 51: 47–60 (1989).

26. Rönnegård L, Valdar W. Detecting major genetic loci controlling phenotypic variability in experimental crosses. Genetics. 188: 435–447 (2011).

27. Bianca Dumitrascu, Gregory Darnell, Julien Ayroles, Barbara E Engelhardt. Statistical tests for detecting variance effects in quantitative trait studies. Bioinformatics. 35: 200–210 (2019).

28. McCaw ZR, Lane JM, Saxena R, Redline S, Lin X. Operating characteristics of the rank-based inverse normal transformation for quantitative trait analysis in genome-wide association studies. Biometrics. 2020 Dec;76(4):1262–72.

29. Gutenbrunner C et al. Tests of linear hypotheses based on regression rank scores. J. Nonparametric Stat., 2: 307–331 (1993).

30. Koenker, Roger. Quantile regression. Vol. 38. Cambridge university press (2005).

31. Song X, Li G, Zhou Z, Wang X, Ionita-Laza I, and Wei Y. (2017). QRank: a novel quantile regression tool for eQTL discovery. Bioinformatics, 33(14), 2123–2130.

32. Liu Y, Xie J. Cauchy combination test: a powerful test with analytic p-value calculation under arbitrary dependency structures. J Am Stat Assoc. 115: 393–402 (2020).

33. Portnoy S, Koenker R. The Gaussian hare and the Laplacian tortoise: computation of squared-error versus absolute-error estimators. Stat Sci. 12: 279–300 (1997).

34. Belloni A Chernozhukov V. l1-penalized quantile regression in high-dimensional sparse models. Ann Stat. 39: 82–130 (2011).

35. Wu Y, Liu Y. Variable selection in quantile regression. Stat Sin. 19: 801–817 (2009).

36. Wang T, Ionita-Laza I, Wei Y. Integrated Quantile RAnk Test (iQRAT) for gene-level associations. The Annals of Applied Statistics. 2022 Sep;16(3):1423–44.

37. Beyerlein A. Quantile regression—opportunities and challenges from a user’s perspective. Am J Epidemiol. 180: 330–331 (2014).

38. Wei Y, Terry MB. Re: “Quantile Regression—Opportunities and Challenges From a User’s Perspective”. American Journal of Epidemiology. 181: 152–153 (2015).

39. Ionita-Laza I, McCallum K, Xu B, Buxbaum JD. A spectral approach integrating functional genomic annotations for coding and noncoding variants. Nat Genet. 48: 214–220 (2016).

40. Kircher M, et al. A general framework for estimating the relative pathogenicity of human genetic variants. Nat Genet. 46: 310–315 (2014).

41. Huang J et al. Genomics and phenomics of body mass index reveals a complex disease network. Nat Commun. 13(1):7973 (2022).

42. Klarin D et al. Genetics of blood lipids among 300,000 multi-ethnic participants of the Million Veteran Program. Nat Genet. 50: 1514–1523. doi: 10.1038/s41588-018-0222-9 (2018).

43. Yang Y, Fan J, Xu H, Fan L, Deng L, Li J, Li D, Li H, Zhang F, Zhao RC. Long noncoding RNA LYPLAL1-AS1 regulates adipogenic differentiation of human mesenchymal stem cells by targeting desmoplakin and inhibiting the Wnt/*β*-catenin pathway. Cell Death Discov. 7(1):105 (2021).

44. Myocardial Infarction Genetics and CARDIoGRAM Exome Consortia Investigators. Coding Variation in ANGPTL4, LPL, and SVEP1 and the Risk of Coronary Disease. N Engl J Med. 374(12):1134–44 (2016).

45. Khera AV et al. Association of Rare and Common Variation in the Lipoprotein Lipase Gene With Coronary Artery Disease. JAMA. 317(9):937–946 (2017).

46. Tcheandjieu C et al. Large-scale genome-wide association study of coronary artery disease in genetically diverse populations. Nat Med. 28(8):1679–1692 (2022).

47. Wojcik GL et al. Genetic analyses of diverse populations improves discovery for complex traits. Nature. 570(7762):514-518 (2019)

48. Kooperberg C, Leblanc M. Increasing the power of identifying gene x gene interactions in genome-wide association studies. Genet Epidemiol. 32: 255–263 (2008)

49. Liis Kolberg, Nurlan Kerimov, Hedi Peterson, Kaur Alasoo. Co-expression analysis reveals interpretable gene modules controlled by trans-acting genetic variants eLife 9:e58705 https://doi.org/10.7554/eLife.58705 (2020).

50. Yun SH, Sim EH, Goh RY, Park JI, Han JY. Platelet Activation: The Mechanisms and Potential Biomarkers. Biomed Res Int. 2016:9060143 (2016).

51. Guo MH, Nandakumar SK, Ulirsch JC, Zekavat SM, Buenrostro JD, Natarajan P, Salem RM, Chiarle R, Mitt M, Kals M, Pärn K, Fischer K, Milani L, Mägi R, Palta P, Gabriel SB, Metspalu A, Lander ES, Kathiresan S, Hirschhorn JN, Esko T, Sankaran VG. Comprehensive population-based genome sequencing provides insight into hematopoietic regulatory mechanisms. Proc Natl Acad Sci U S A. 114(3):E327–E336. doi: 10.1073/pnas.1619052114. (2017)

52. Zhang X, Johnson AD, Hendricks AE, Hwang SJ, Tanriverdi K, Ganesh SK, Smith NL, Peyser PA, Freedman JE, O’Donnell CJ. Genetic associations with expression for genes implicated in GWAS studies for atherosclerotic cardiovascular disease and blood phenotypes. Hum Mol Genet. 23: 782–795. doi: 10.1093/hmg/ddt461 (2014)

53. Geraci M. Linear Quantile Mixed Models: The lqmm Package for Laplace Quantile Regression. Journal of Statistical Software, 57: 1–29 (2014).

54. Geraci M, Bottai M. Linear Quantile Mixed Models. Statistics and Computing, 24: 461– 479 (2014).

55. Kong A, Thorleifsson G, Frigge ML, Vilhjalmsson BJ, Young AI, Thorgeirsson TE, Benonisdottir S, Oddsson A, Halldorsson BV, Masson G, Gudbjartsson DF, Helgason A, Bjornsdottir G, Thorsteinsdottir U, Stefansson K. The nature of nurture: Effects of parental genotypes. Science. 359: 424–428 (2018).

56. Chang, C. C., et al. Second-generation PLINK: rising to the challenge of larger and richer datasets. Gigascience 4, 7 (2015).

57. Sinnott-Armstrong, N, et al. Genetics of 35 blood and urine biomarkers in the UK Biobank. Nat Genet. 53: 185–194 (2021).

58. Manichaikul, A. et al. Robust relationship inference in genome-wide association studies. Bioinformatics. 26: 2867–2873 (2010).

